# Trade-offs and design principles in the spatial organization of catalytic particles

**DOI:** 10.1101/2020.06.14.146076

**Authors:** Florian Hinzpeter, Filipe Tostevin, Alexander Buchner, Ulrich Gerland

## Abstract

Spatial organization of catalytic particles is ubiquitous in biological systems across different length scales, from enzyme complexes to metabolically coupled cells. Despite the different scales, these systems share common features of localized reactions with partially hindered diffusive transport, determined by the collective arrangement of the catalysts. Yet it remains largely unexplored how different arrangements affect the interplay between the reaction and transport dynamics, which ultimately determines the flux through the reaction pathway. Here we show that two fundamental trade-offs arise, the first between efficient inter-catalyst transport and depletion of substrate, and the second between steric confinement of intermediate products and accessibility of catalysts to substrate. We use a model reaction pathway to characterize the general design principles for the arrangement of catalysts that emerge from the interplay of these trade-offs. We find that the question of optimal catalyst arrangements generalizes the famous Thomson problem of electrostatics.

---

The physics underlying the spatial organization of particles in dense systems has a long history and displays intriguing behaviors [1]. Traditionally, physics has dealt only with inert particles, while catalytic particles are key to the complexity of chemical and biological systems. The physics of spatially arranged catalytic particles remains largely unexplored. Biology provides many examples of systems where particles that catalyze sequential reactions exhibit a striking degree of spatial organization across all length-scales of living systems [2], from individual molecules to collections of cells.

At the molecular scale, enzymes that catalyze consecutive reactions in a biochemical pathway are often organized into multi-enzyme complexes, micro-compartments, or other assemblages [3–6]. Well known examples include cellulosomes [7], purinosomes [8], and carboxysomes [9].

Such arrangements are thought to enable increased efficiencies and reaction yields by keeping metabolic intermediates between enzymatic steps out of equilibrium with the bulk solution, a concept referred to as ‘substrate channeling’ [10]. Recently, a variety of scaffolding and confinement approaches were leveraged in efforts to design efficient spatially-organized multi-enzyme reactions *in vitro* [11–13]. Similar ideas of efficiently arranging consecutive catalysts are pursued in the realm of concurrent tandem catalysis to improve the yield and specificity of sequential chemical reactions [14–16]. Understanding the underlying physical principles is crucial for engineering such efficient multi-catalyst systems. For instance, it remains controversial whether proximity of consecutive enzymes alone is sufficient to allow channeling of diffusing intermediates [17–20], or whether additional confinement of intermediates or active mechanisms are required [10].

At a higher level of organization, enzyme compartments and complexes themselves can be seen as catalytic particles. These superstructures also function synergistically and some have been found to spatially colocalize [21–23]. For example, purinosomes in HeLa cells were found to localize to mitochondria [23]. Their spatial proximity appears to ensure that mitochondrial-derived metabolic intermediates are efficiently captured by purinosomes to enhance nucleotide production [23, 24].

On an even larger scale, whole cells can be considered as catalytic particles. By taking up, processing, and secreting biochemical molecules, cells effectively function as catalysts that alter the chemical composition of their environment. Notably, on this multicellular scale, different cells also work together to sequentially process metabolites [25–28]. For example, biological nitrification, the conversion of ammonia to nitrate via the intermediate nitrite, is performed by two specialized microbes [29]. The first step, the oxidation of ammonia to nitrite, is catalyzed by ammonia-oxidizing microbes, while the second step, the conversion of nitrite to nitrate, is performed by nitrite-oxidizing bacteria. These synergistic microbes grow together in spatially structured biofilms [30].

Despite the differences in length scale, the behavior of these systems is often governed by common underlying physics. The metabolites are typically small molecules that move freely by diffusion, while the catalysts are typically much larger and are spatially localized or constrained in their motion relative to each other. The reaction fluxes are determined by a kinetic interplay between diffusive transport of metabolites and the reaction kinetics at the specific locations of the catalysts. Previously, the reaction-diffusion dynamics of spatially arranged catalysts were studied with continuum models, which do not account for the discrete nature of catalysts but describe their arrangement by density profiles on mesoscopic length scales [31–33]. This prior work analyzed the impact of the large-scale catalyst arrangement on the overall reaction efficiency. It is currently unclear how the discrete nature of the catalysts affects this efficiency. If multiple catalysts are placed in close proximity, as in enzyme clusters or microbial biofilms, the resulting “crowded” geometries lead to a complex spatial network of diffusive fluxes that exchange the participating metabolites between the catalysts and with the environment. Previous studies characterized the effects of “random crowding” on the diffusion and reaction dynamics of molecules [34–38]. In contrast, the possibility for “designed crowding”, in which arrangements of objects are chosen such as to selectively block or direct the diffusion of molecules, and simultaneously to catalyze their biochemical conversion, remains largely unexplored. When and how should the arrangement of individual catalysts be tuned such as to promote reactions along a reaction pathway? Which trade-offs and design principles emerge from the interplay of the physical processes described above?

Here we compare different strategies for arranging catalysts, using a model of discrete catalysts together with continuous reaction-diffusion dynamics for metabolites. Our model is not designed to describe the detailed properties of specific catalysts, but to identify general physical principles that apply to all systems of this type. We identify spatial organization strategies that are advantageous in different parameter regimes of catalyst reactivity and metabolite diffusion. We find that in the reaction-limited regime, where the catalytic reaction is slow compared to diffusion, the best strategy is to colocalize the catalysts into large clusters. In contrast, in the diffusion-limited regime it is beneficial to form pairs or small complexes of catalysts. The enhancement of the reaction flux compared to unordered, delocalized catalyst arrangements is highest when the catalyst concentrations are low. The change of the optimal localization strategy arises from two trade-offs: First, a compromise between efficient transfer of intermediates and competition for substrates. Second, a tradeoff between steric shielding and confinement of metabolites. The interplay of these effects gives rise to non-trivial symmetries of the optimal configurations of model multi-catalyst complexes.

## MODEL

We consider a model two-step catalytic reaction performed by two catalysts (Fig. 1a). The first catalyst 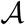 converts a substrate 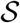 to an intermediate product 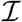, which is subsequently converted to the product 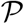 by the second catalyst 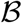,

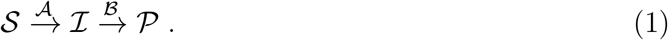

**FIG. 1:**
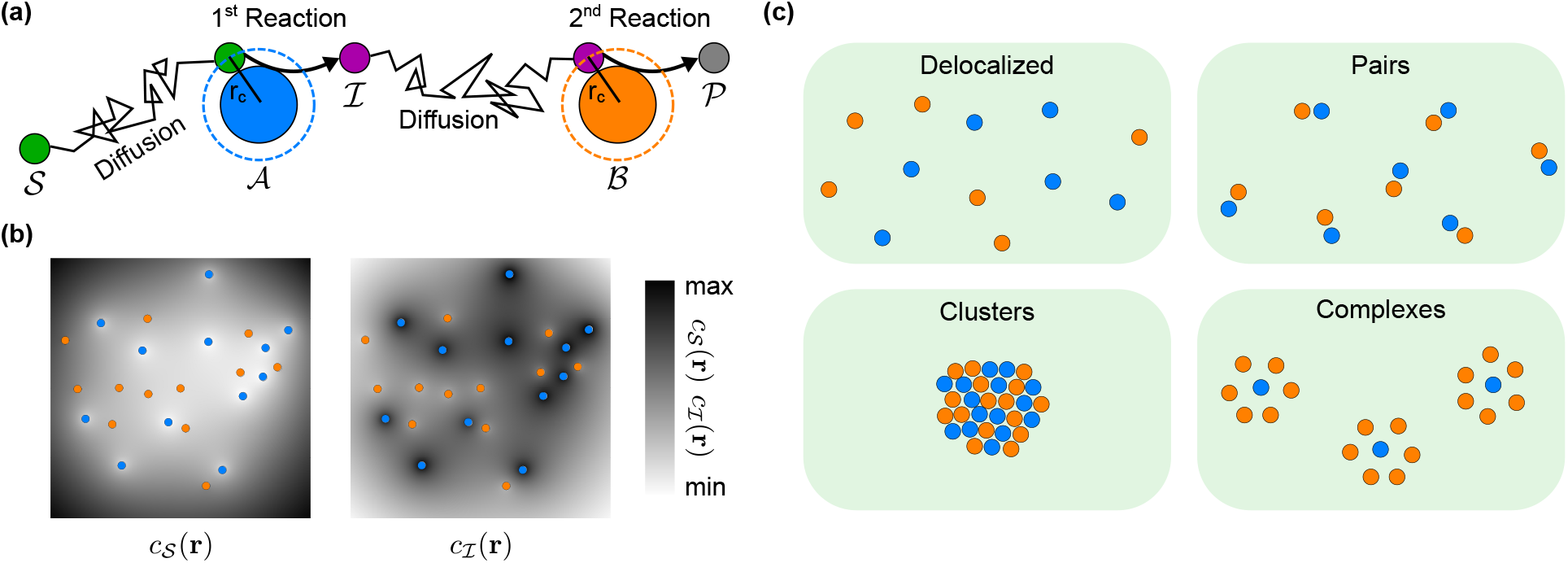
Schematic illustration of the model. (a) A model two-step reaction pathway involving the catalysts 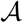 and 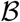, the substrate 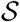, the intermediate product 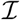, and the product 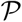. The metabolites 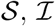 diffuse and react when they come into contact with the catalysts 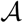 and 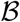, respectively (contact distance *r_c_*). (b) For a given spatial arrangement of the catalysts (blue and orange disks), the reaction-diffusion dynamics of the metabolites 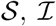 is described in terms of continuous concentration profiles (the steady-state profiles, 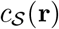 and 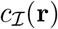, are shown as grayscale gradients). This description ignores steric exclusion between metabolites, but incorporates steric exclusion of metabolites by catalysts via the effective radius rc of the spheres that represent the catalysts. (c) Examples of catalyst arrangements analyzed here: Random delocalized arrangements, 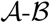 pairs, high-density clusters of randomly arranged catalysts, and complexes with a fixed stoichiometry and geometry.

Between catalysts, metabolites move by diffusion. In general, the catalysts can be syntrophic microbes, metabolically linked organelles in eukaryotic cells or consecutive enzymes of the same biochemical pathways, while metabolites are typically small molecules. We model the catalysts as spatially-extended spherical particles with an effective interaction radius *r_c_ = r_cat_* + *r_met_*, while treating the metabolites 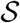 and 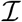 as point like particles and describing their distributions as continuous concentration profiles, 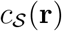 and 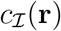 (see Fig. 1b and ‘Methods’). This description is adequate as long as metabolites do not cause significant steric hindrance to one another over the length scale of catalysts, such as when metabolites are significantly smaller than catalysts. This is typically the case. For example, the sizes of the commonly-studied sequential enzymes glucose oxidase (GOx) and horseradish peroxidase (HRP) can be approximated by their hydrodynamic radii of ~43Å [39] and ~30Å [40], respectively, while the sizes of their substrate metabolites glucose and H_2_O_2_ are approximately ~4Å [41] and ~3Å [42]. For molecular-scale metabolites moving between micrometer-sized organelles or cells, the separation of sizes is even larger.

The reaction pathway is supplied with substrate 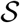 from the surrounding environment, where we assume a fixed level of 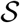. Intermediates 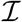 can escape the reaction to the environment where their concentration is negligible. The behavior of this model depends on two dimensionless parameters 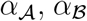, which capture the relative timescales of reactions and diffusion for each catalyst. Both can be expressed in the form

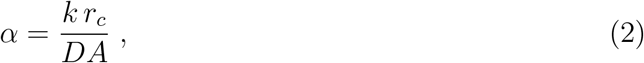

where *D* is the respective metabolite diffusion coefficient, *k* is the intrinsic catalytic efficiency of the catalyst, *A* is the surface area of the catalyst, and *r_c_* the interaction radius (see ‘Methods’). The intrinsic catalytic efficiency *k* differs from the macroscopic catalytic efficiency, *k_cat_/K_M_*, since the latter incorporates the effective timescale of metabolite-catalyst encounters due to diffusion, whereas we describe diffusion explicitly. Hence *k* is an effective catalytic efficiency for metabolites “at contact” with the catalyst, which captures molecular-scale details of the metabolite-catalyst interaction.

Experimentally relevant values of the reaction-diffusion parameter *α* span a wide range. For substrates with diffusion coefficient *D* ≈ 100 μm^2^ s^-1^, a reaction-limited enzyme with catalytic efficiency *k* ~ 10^6^ M^-1^ s^-1^ and r_c_ ≈ 2nm has *α* ~ 10^-3^. For fast, diffusion-limited enzymes, the observed macroscopic catalytic efficiency *k*_cat_/*K*_M_ is around 10^9^ M^-1^ s^-1^, while the intrinsic catalytic efficiency *k* can be 1-2 orders of magnitude higher (see ‘Methods’), implying that *α* reaches values of up to 100. When the catalysts are synergistic microorganisms, we also expect the *α* values to fall within this range, e.g. for the uptake of ammonia by *N. maritimus*, which functions as an ammonia oxidizer in two-step nitrification (*k*_cat_/*K*_M_ ‘ 6.6 × 10^11^ M^-1^ s^-1^ [43], *r_c_* = 0.4 μm, *D* = 1000 μm^2^ s^-1^), we obtain *α* ~ 0.2. Given these estimates, and our aim to broadly explore the physical principles of spatially organized catalytic particles, we characterize the behavior of our model over the entire range of *α* values.

All steady-state properties of our model systems depend only on the reaction-diffusion parameters 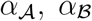, the number of catalysts, 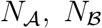 of each type, and their spatial arrangement. We investigate how different strategies for arranging the catalysts affect the steady-state pathway flux

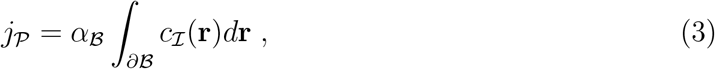

where 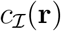 is the steady-state concentration profile of intermediates, the integral is taken over the surfaces of all 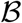 catalysts, and the timescale has been rescaled by 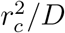 (see ‘Methods’).

It is important to note that without an alternative mechanism by which intermediates can be lost from the system, the steady-state rate of product formation must equal the rate of substrate consumption, irrespective of the spatial organization of the catalysts. Intermediates can be lost through various mechanisms such as leakage through a permeable system boundary, consumption in competing reactions, or spontaneous decay if the intermediate is intrinsically unstable. We consider all of these possibilities, but use the case where intermediates escape upon reaching the system boundary to illustrate the system behavior in the main text. To obtain a comprehensive understanding that applies to different biological scenarios like bacterial biofilms on a surface or clusters of enzymes in the cellular cytoplasm, we study systems in both two and three dimensions (“2D” vs. “3D”).

## RESULTS

### Randomly positioned catalysts

We first analyze ensembles of random catalyst arrangements, as a reference to compare against specific localization strategies. These random ensembles also allow us to identify characteristics of the catalyst arrangements that correlate with the pathway flux (3). We considered a range of different values for the reaction-diffusion parameters and catalyst abundances, which for simplicity we chose symmetrically (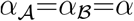 and 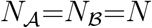). For each parameter set we computed the pathway fluxes 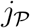 for 3000 random catalyst arrangements (see ‘Methods’). We limited these arrangements to a 2D geometry, where the smaller configuration space allows for a more thorough sampling of catalyst arrangements. We study the behavior of 3D systems further below, when we analyze specific localization strategies.

Figure 2a (top) shows the behavior of the mean pathway flux 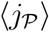, i.e. the ensemble average over all catalyst configurations. For small reaction-diffusion parameters, the mean flux increases quadratically with a, while it saturates for *α* ≫ 10. This behavior reflects the transition of the system from a reaction-limited regime, in which each reaction rate is limited by the probability *p ~ α* ≪ 1 that a metabolite-catalyst encounter results in a reaction (with the quadratic increase coming from the fact that the pathway consists of two reactions that each scale with *α*), to a diffusion-limited regime at large a, in which most encounters are reactive, *p* ≈ 1, and the rate of reactions is instead set by the frequency of such encounters due to diffusion.

**FIG. 2:**
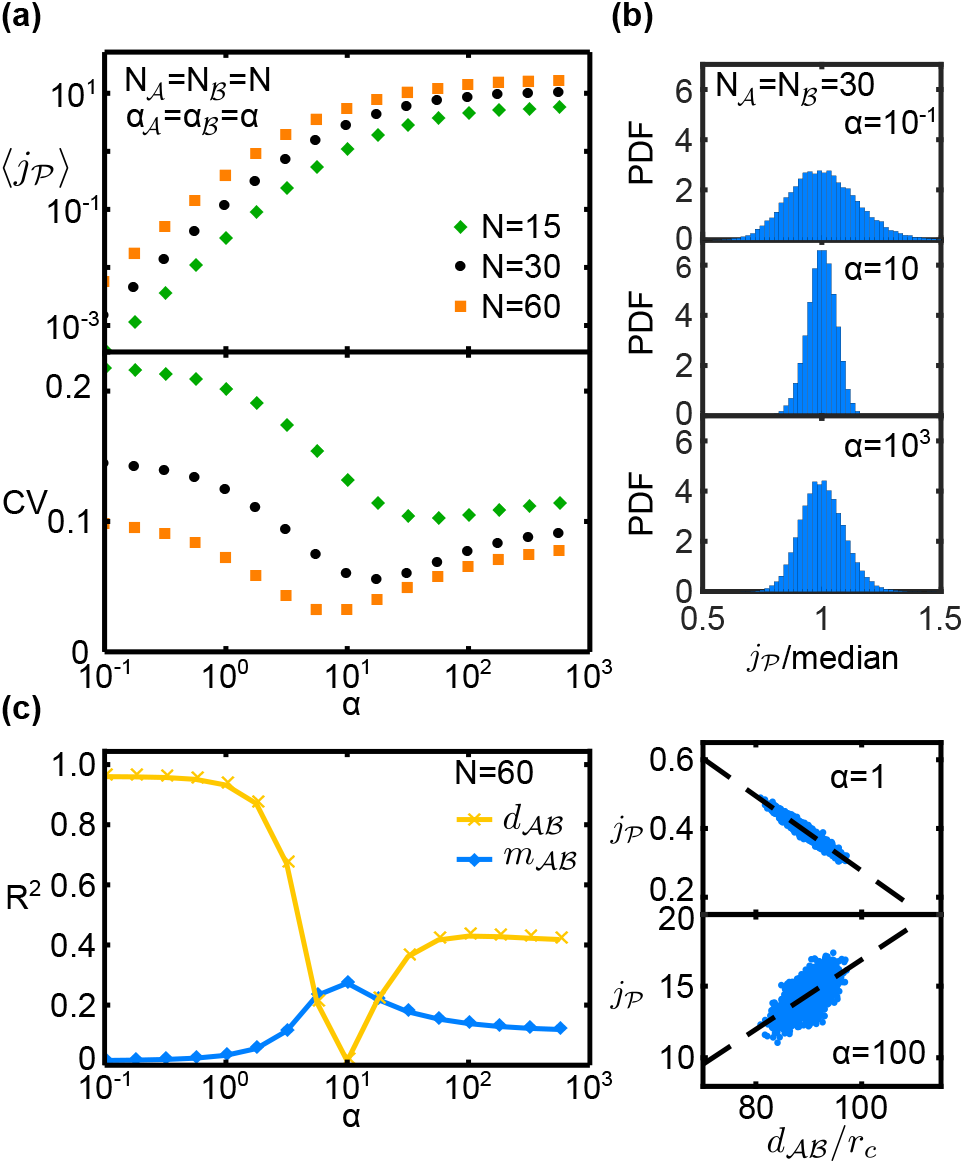
Properties of the pathway flux 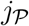 for randomly positioned catalysts (two-dimensional system with absorbing boundary for intermediates, parameters chosen symmetrically with 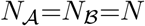 and 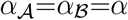). (a) The mean flux 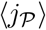 (top) and coefficient of variation *CV* (bottom) as *α* is varied for different values of *N*. (b) Histogram of reaction fluxes for *N* = 30 at three different values of *α*. (c) Coefficient of determination *R*^2^ of the linear regression of 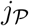 against 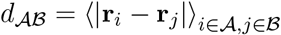 (crosses) and 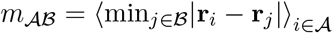 (diamonds), and regression plots against 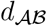 at low and high values of *α*.

### Correlations between catalyst arrangements and pathway fluxes

To examine how sensitive the pathway flux is to the catalyst arrangement, we inspected the distribution of all flux values 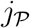 (at given *α* and *N*). Fig. 2b shows that the distribution is significantly narrower around *α* ≈ 10 than at smaller or larger *α* values (at fixed *N* = 30). The variability of the flux, as measured by the coefficient of variation, 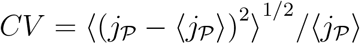, indeed displays a non-monotonic dependence on the reactiondiffusion parameter *α*, with a minimum within the crossover region from the reaction-limited to the diffusion-limited regime (Fig. 2a, bottom). At this point, the configuration of the catalysts has the least impact on the pathway flux. On the other hand, at small or large *α* the variation is larger, implying that the reaction flux is more sensitive to the particular arrangement of catalysts.

We next sought properties of the catalyst arrangements that correlate with changes in 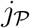 in different *α* regimes. For small *α* (reaction-limited regime), the pathway flux of a given configuration is strongly anti-correlated with the mean distance between 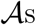 and 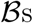, 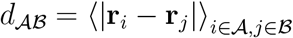 (*R*^2^ > 0.95, Fig. 2c). For large *α* (diffusion-limited regime), there is a weaker (*R*^2^ ≈ 0.4) but positive correlation between these quantities. However, in the transition region, the correlation is insignificant (*R*^2^ ≈ 0.01). Similar trends, albeit with weaker correlations, are also observed for the distances 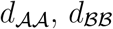, and the mean radial coordinates 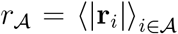 and 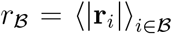 (Fig. S1). While these properties all displayed negligible correlations with the flux in the transition region, we observed a significant positive correlation (*R*^2^ ≈ 0.27) between the flux and the minimal distance between 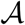 and 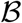 catalysts, 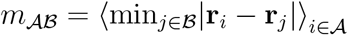 in this region (Fig. 2c).

Together these data indicate that for small *α* the highest flux is generated by configurations in which the catalysts are generally placed closer together, and closer to the center of the system. In contrast, for large *α* the highest flux comes from configurations where the reaction centers are further apart, and closer to the periphery of the system. At the point where the variability in the flux is smallest, both the best- and worst-performing configurations have similar mean separations. However there remains an advantage to placing catalysts such that each is in close proximity to at least one reaction center of the other type.

### Comparison of different localization strategies

Having seen that the pathway flux depends on such quantities as the average and minimal distance between catalysts, we investigated in more detail two extreme localization strategies that emphasize these properties (Fig. 1c): (i) a single dense but disordered cluster of catalysts, and (ii) pairs of catalysts consisting of one 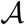 and one 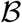 at a separation of rc. We again consider mean fluxes 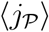, averaged over an ensemble of configurations where the catalysts are either paired or clustered (see ‘Methods’). Figures 3a and 3b show 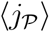 as a function of *α* in two and three dimensions, respectively, for pairs, clusters, as well as our reference case of random arrangements (‘delocalized’).

**FIG. 3:**
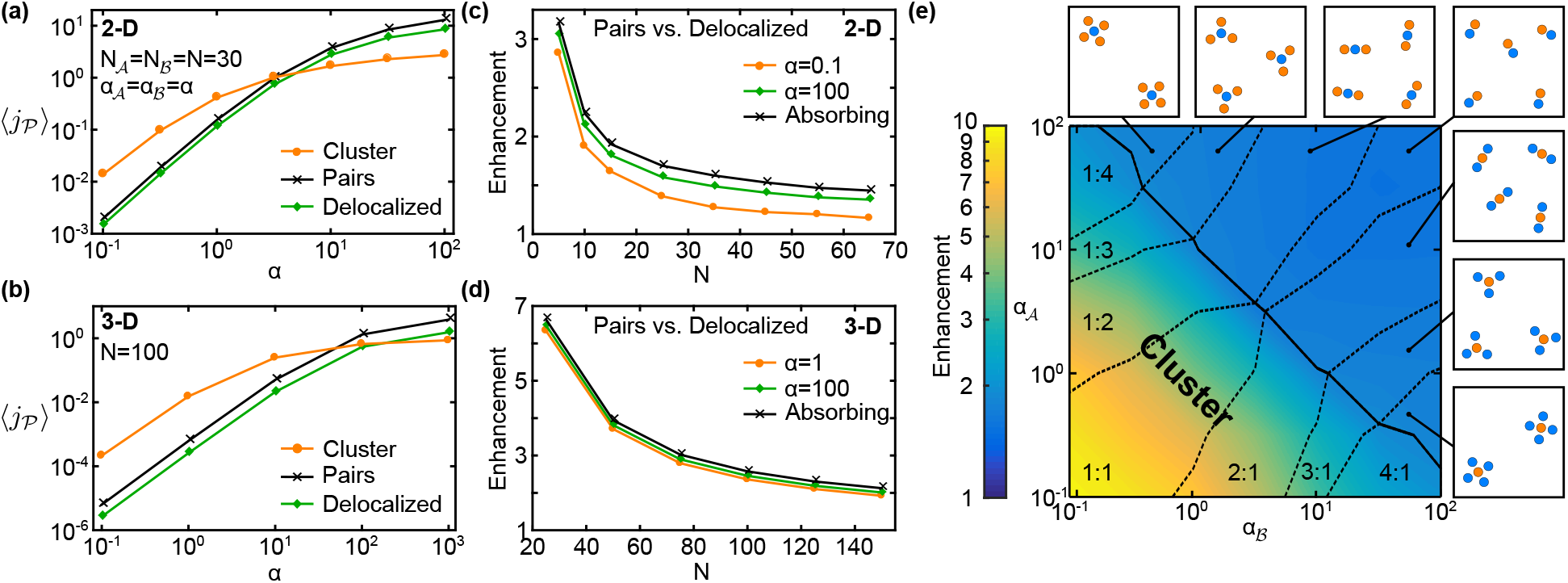
Comparison of different localization strategies. Mean pathway fluxes of clustered arrangements, 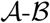-pair arrangements, and completely random arrangements (‘delocalized’) in (a) two-dimensional and (b) three-dimensional systems (parameter values as indicated). Intermediates can be lost via the absorbing system boundary. The corresponding enhancement of the mean flux in pair arrangements relative to delocalized arrangements as the catalyst number is varied is shown in (c) and (d), respectively. (e) Phase diagram of optimal stoichiometries and spatial organization of catalysts in 2D, as 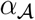 and 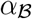 are varied given a constant total number of catalysts, 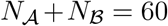. The solid line separates the regime where clustering is the optimal strategy from that in which small complexes produce a higher flux; dashed lines denote changes in the optimal ratio 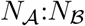. The color scale represents the enhancement of the mean flux relative to delocalized arrangements.

We observe that the mean flux for pair arrangements is always larger than for delocalized arrangements. This reflects the fact that placing 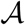 and 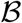 in close proximity increases the probability that a produced 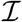 molecule will encounter at least one 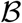 and react before diffusing to the boundary of the system. The flux enhancement is largest at low catalyst concentrations (small *N*) and in three dimensions (Fig. 3c and d), reaching almost 7-fold for *N* = 20, which for an enzymatic system corresponds to a catalyst concentration of ~1 μM. This lies within the range of typical intracellular enzyme concentrations spanning from high nanomolar to micromolar concentrations [44]. The observed dependence of the enhancement on the catalyst concentration is in line with the diffusive capture probability: In random arrangements, a low concentration implies a large mean separation between consecutive catalysts, such that 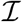 molecules are unlikely to encounter a 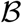 before diffusing out of the system. At higher concentrations, catalysts are already closeby when randomly arranged, such that 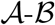-pair formation barely increases the capture probability.

The effect of the localization strategy on the pathway flux could also depend on the mechanism of intermediate loss and the associated loss rate. To characterize this dependence, we considered modified models where intermediates either leak out through the system boundary at a reduced rate (Figs. S2 and S3), or are consumed within the system by an alternative mechanism (Fig. S4). The flux enhancement due to 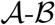-pair formation (Fig. 3c) is reduced if intermediates leak out of the system at a limited rate, such that the boundary is only partially absorbing (Fig. S3). This is because for small leakage rates, intermediates can explore a larger fraction of the system and thereby come in contact with more 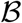 catalysts. This effect attenuates the beneficial effect of catalyst proximity, which relies on the rapid transfer of intermediates from one catalyst to the next. The behavior for intermediate loss within the system, either via a competing pathway or a decay reaction, is qualitatively similar (Fig. S4).

The flux enhancement obtained by the pair strategy can also be described by an approximate analytical expression (Fig. S5), which shows how the enhancement depends on the concentration of catalyst pairs, the distance between the paired catalysts, and the average distance an intermediate diffuses before it is lost (Fig. S6). Notably, the enhancement is independent of the catalytic efficiencies of the catalysts. This description rationalizes why no measurable enhancement could be detected in the experiment of Ref. [20], see Fig. S7.

Turning now to the clustering strategy, we found that for small *α* the clustered configurations achieve a significantly higher mean flux than the delocalized and pair arrangements. This enhancement is approximately ten-fold in 2D and hundred-fold in 3D for similar numbers of catalysts (Fig. 3a and b). However, as *α* is increased, a transition occurs into a regime where the pair arrangements produce a higher mean flux than the clusters. When *α* is further increased, even the delocalized arrangements outperform the cluster. The quantitative characteristics of this behavior also depend on the loss mechanism and loss rate (Figs. S2 and S4). Furthermore, nonlinear reaction kinetics due to catalyst saturation can affect the relative performance of the different spatial strategies (Figs. S8 and S9). However, the behavior is robust in its qualitative features. In particular, the transition between the clustering and the pairing strategies exists as long as the catalysts are not fully saturated.

To generalize these observations we considered asymmetric systems where the 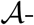 and 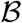-catalysts have different reaction-diffusion parameters 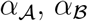 and copy numbers, 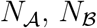. Figure 3e summarizes the results in the form of a phase diagram showing the 2D configuration that produces the highest mean pathway flux with a given total catalyst number, 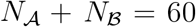, but different 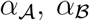. In the region defined approximately by 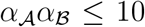 (solid line), the highest flux was produced by a single large cluster. In contrast, for larger 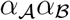, small complexes produced a higher flux. In both regimes, the relative values of 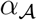 and 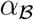 determined the optimal catalyst stoichiometry, favoring a larger number of 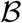 than 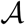 in regions with 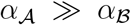, but more 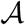 than 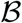 when 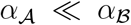. Interestingly, despite the difference in boundary conditions for 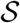 and 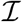, the phase diagram is approximately symmetrical about the line 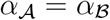.

The transition in the optimal arrangement, from a cluster at small *α* to a more disperse arrangement at large *α* is reminiscent of a previously reported transition in the optimal density profile of enzymes around a localized source [31, 32]. However, those studies did not incorporate several physical effects considered here, in particular the impact of the catalyst arrangement on the first reaction flux, as well as steric effects due to the discreteness of catalysts. How does the interplay between the different effects result in the behavior observed in Fig. 3?

### Trade-off between substrate depletion and efficient intermediate transfer

To disentangle the different effects contributing to the performance of spatially organized catalysts, we eliminated all effects caused by steric exclusion, by allowing metabolites to diffuse through the space occupied by catalysts, and reactions to occur throughout their volume rather than on their surface (see Supplementary Information). This modified model displays the same qualitative behavior as our full model with respect to the comparison of different localization strategies (Fig. S10), implying that the transition in Figs. 3a,b results from a trade-off between substrate depletion and efficient intermediate transfer: For small *α*, intermediates are likely to escape without reacting even if the catalysts are arranged in pairs, since the reaction probability at each 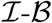 encounter is low. This loss is attenuated by clustering several copies of 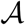 and 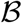. An 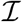 molecule produced in such a cluster has a higher probability to be processed by a proximal 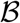, even if the probability of reaction with each individual 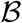 is low [33]. As *α* becomes larger the reaction probability at each 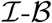 encounter increases, reducing the benefit of clustering. Furthermore, since each 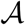 consumes more of the incoming 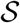, steeper concentration gradients develop around the cluster. This substrate depletion reduces the productivity of 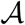’s in the cluster as they effectively compete for substrate. It then becomes increasingly unfavorable to position 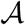’s into close proximity of one another [45–47]. Instead, a more disperse arrangement, with larger distances between catalysts of the same type, becomes preferable.

A quantitative comparison between the full model and the modified model reveals the steric exclusion effects on the relative performance of different localization strategies. In the case of delocalized catalysts, we found little difference in the flux between the model with permeable catalysts and the full model with impermeable catalysts (Fig. S11). However, for clustered catalyst configurations the pathway fluxes of the two models differ significantly (Fig. 4a,b). Especially in the reaction-limited regime (small a) the impermeability of reaction centers significantly enhances the pathway flux. This enhancement increases with the packing density *ϕ* of catalysts in the cluster and weakly with the number of catalysts, reaching almost five-fold in 2D for *ϕ* = 0.8 and 1.7-fold in 3D for *ϕ* = 0.6 (Fig. 4c,d top). In the diffusion-limited regime (large a), on the other hand, we find that the impermeability of catalysts leads to a reduction of the pathway flux, which is strongest for high packing densities and large numbers of catalysts (Fig. 4c,d bottom). This shows that steric exclusion reinforces the trade-off between substrate depletion and intermediate exchange, by decreasing the flux in the reaction-limited regime and increasing the flux in the diffusion-limited regime.

**FIG. 4:**
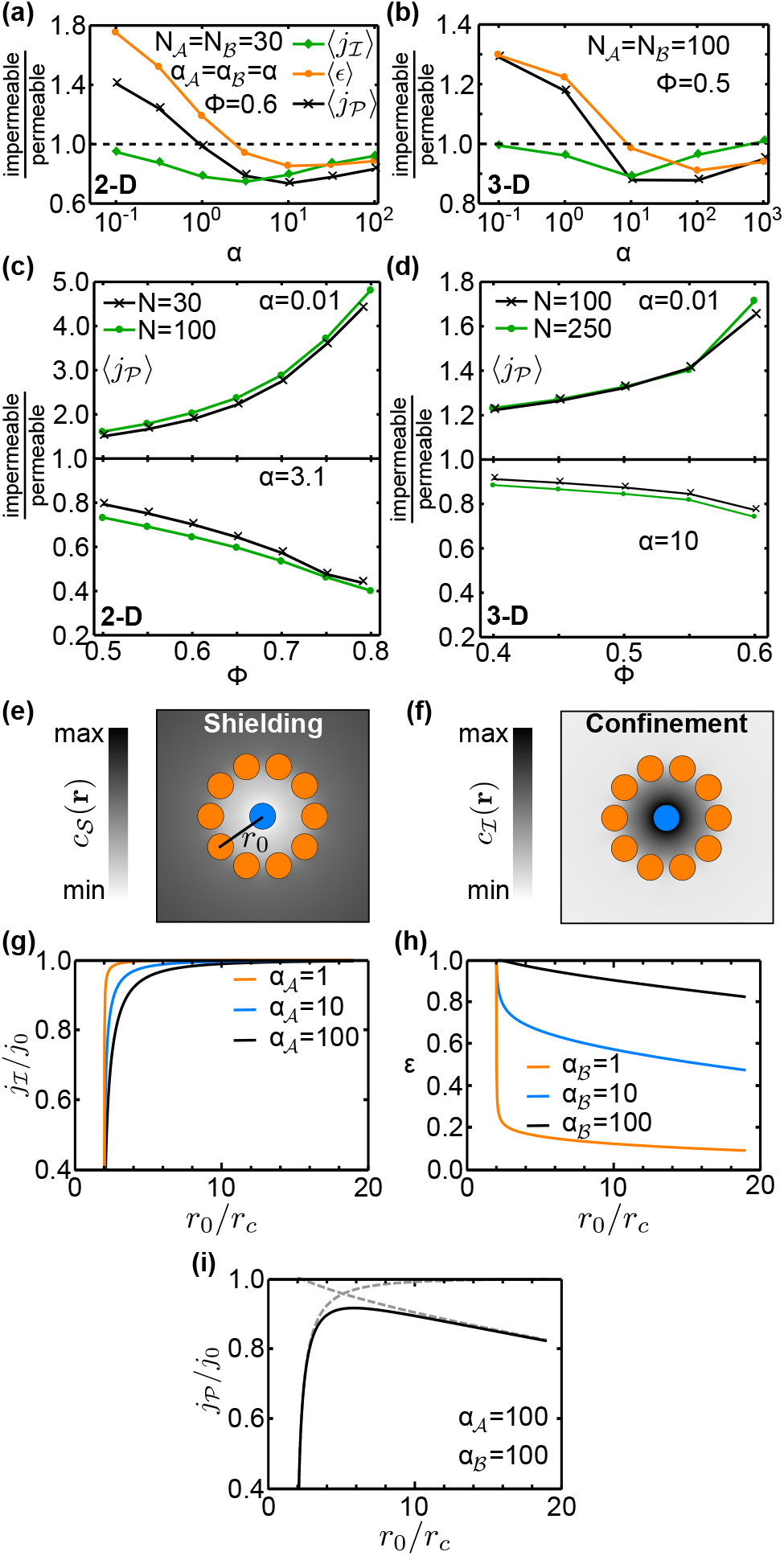
Trade-off between substrate shielding and confinement of intermediates. (a,b) Steric exclusion effects for clustered catalyst arrangements, characterized by the ratios (impermeable to permeable) of the mean fluxes 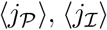, and the mean efficiency 〈*ϵ*〉 for (a) 2D systems with packing density *ϕ*=0.6 and (b) 3D systems with *ϕ*=0.5 (lines are guides to the eye). (c,d) Dependence of the steric exclusion effect in 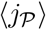 on the packing density in (c) 2D and (d) 3D. (e) Illustration of substrate shielding in a catalyst arrangement of a central 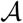 surrounded by a ring of ten 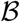. Diffusion of substrate 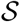 to 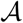 is hindered, causing a reduced concentration 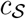 inside the ring. (f) Conversely, intermediates 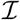 produced by 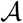 do not easily diffuse out, increasing 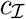 within the ring (confinement). (g,h) The effect of substrate shielding is reflected in (g) the dependence of the flux 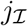 on the ring radius *r*_0_, while the confinementeffect is reflected in (h) the conversion efficiency *ϵ*. (i) Together, these effects produce a non-monotonic pathway flux, with an optimal *r*_0_ where 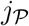 is maximal, demonstrating the trade-off between 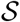 shielding and 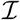 confinement. All fluxes are plotted relative to *j*_0_, the value of 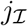 in the absence of a 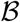 ring.

### Trade-off between metabolite shielding and confinement

For a more comprehensive understanding of trade-offs in the spatial organization of sequential catalysts, we analyze the pathway flux 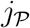 together with the flux 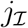 of the first reaction (defined in Eq. 10, analogously to 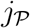). The efficiency 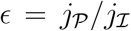 of the second reaction correponds to the fraction of produced 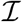 converted to 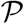. Figures 4a,b show that the mean flux 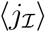 is reduced across the full *α* range when catalysts obstruct diffusion in the clusters, but this reduction is strongest at intermediate *α* values. In contrast, the mean efficiency 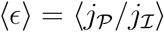 displays the same qualitative behavior as 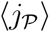 but with larger amplitude. These behaviors arise from the interplay of two steric exclusion effects within clusters, the “shielding” of catalysts from metabolites, and the “confinement” (diffusive trapping) of intermediates.

To illustrate these effects, we consider the special arrangement of several 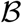 on a ring of radius *r*_0_ around a central 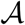. Such an arrangement could approximate the environment around a single 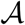 in a cluster, and allows us to monitor how the flux of each reaction varies as a function of 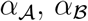, and the clustering density, which we set via *r*_0_. As *r*_0_ is decreased, the 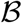 ring progressively blocks the diffusion of 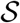 into the vicinity of the 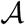 catalyst, leading to a marked reduction in 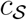 (Fig. 4e). This shielding of 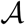 from its substrate decreases the intermediate flux 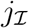 (Fig. 4g). The magnitude of the 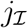 reduction, and the radius at which 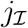 starts to decrease, both increase with 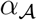 For the second reaction, the presence of the 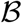 ring restricts the diffusion of 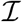 out of the ring, increasing the local concentration of 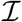 within the ring (Fig. 4f) and therefore the conversion efficiency of 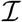 into 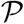 (Fig. 4h). This confinement effect is largest at small 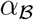, when the probability of reaction in each 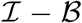 encounter is lowest. The trade-off between shielding of substrate and confinement of 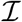 leads to the emergence of an optimal ring radius at the point where the decline of 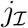 due to shielding is exactly balanced by the increased efficiency of 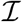-processing achieved by the confinement (Fig. 4i).

Returning now to the scenario of a dense cluster containing both 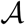 and 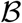, we conclude from Figs. 4a,b that shielding tends to reduce 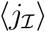 most significantly at intermediate values of *α*. When *α* is small, reactions are slow and 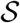 molecules nevertheless have sufficient time to diffuse throughout the cluster. At the opposite extreme of large *α* and fast reactions, 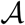 catalysts on the periphery of the cluster are effectively able to consume most of the available 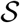, such that little substrate reaches the center of the cluster even when diffusion is unimpeded.

For the second reaction, the confinement of 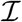 that is produced within the cluster increases the conversion efficiency *ϵ* predominantly at small *α*. In the large-*α* regime, however, shielding also dominates and reduces the efficiency of the second reaction. Here, since 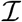 is produced primarily at the periphery of the cluster, it is effectively shielded from 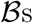 within the cluster, increasing the chance of it diffusing out of the system rather than reacting.

### Geometries of optimal complexes

When catalysts are not randomly clustered, but arranged into complexes with fixed stoichiometry and geometry (Fig. 1c), the question arises: which arrangement is optimal? Experimentally, this question is raised by synthetic biology efforts that use various scaffolding strategies to construct efficient multi-enzyme complexes [48–50], inspired by natural complexes consisting of multiple enzymes in intricate arrangements [51]. Here, we explore this question from a theoretical perspective, starting with the idealized scenario of spherical catalysts that can be freely arranged into complexes with any geometry: how should the catalysts be arranged to globally maximize the pathway flux 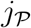? As we found above for ring arrangements, we expect that the optimal geometry will arise as a compromise between the advantageous effects of proximity and confinement and the detrimental effects of shielding and intermediate depletion. Since the relative magnitudes of these effects depend on the reaction-diffusion parameters 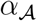 and 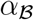, the optimal complex geometry should also depend on these parameters.

For complexes of a single 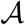 catalyst surrounded by several 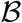 catalysts, the numerically determined optimal geometries (see ‘Methods’) are shown in Fig. 5, for different 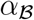 values with 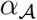 fixed. Surprisingly, the symmetries of the optimal configurations change not only with 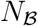, but also as a function of 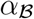. In 2D with 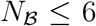 it is always optimal to arrange the 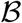 at equidistant positions on a ring around 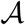 (point symmetry group 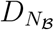); ring radius increases with 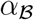 as described above). In contrast, for 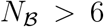, the optimal complexes take on more intricate geometries. At small 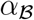 values, the complexes leave tight channels open for substrate to enter, but intermediates unlikely to escape without making multiple contacts with 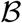 catalysts. For larger 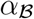, the 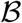 are typically divided between an inner ring and an outer population arranged at angular positions corresponding to the gaps in the inner ring. For even 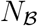, these arrangements are concentric rings that are rotated by 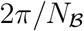 with respect to one another (point group 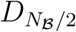). For odd 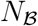, it is not possible to form two full rings and thus the arrangement shows only a single reflection symmetry axis. Interestingly, in the limit of extremely large 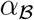, the optimal arrangement changes from these star-like arrangements back to a single ring (point group 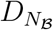), provided 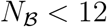.

**FIG. 5:**
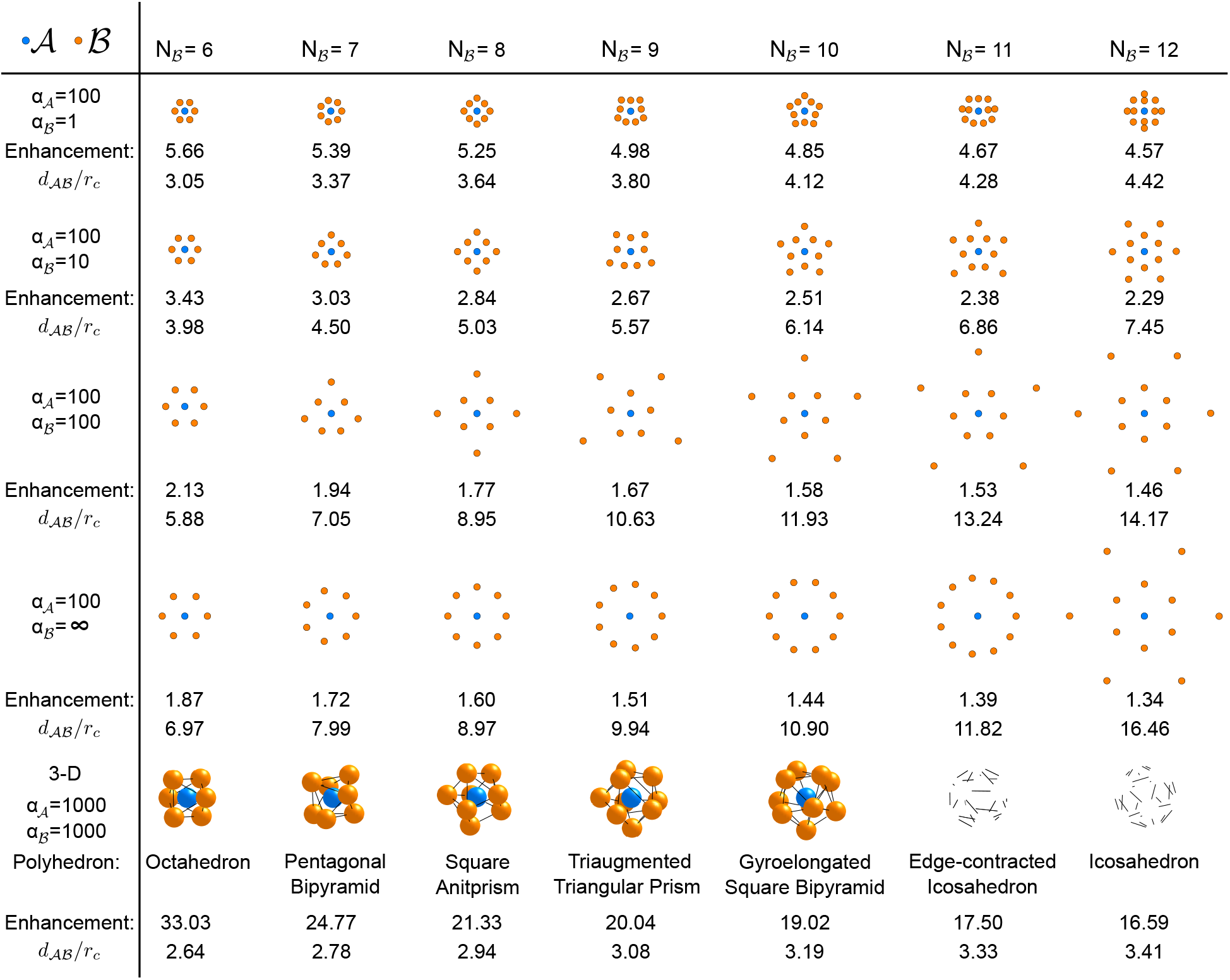
Optimal geometries of complexes with a single 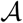 catalyst (blue) surrounded by different numbers 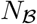 of 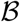 catalysts (orange), in 2D and 3D. For the 2D case, it is shown how the optimal geometry changes as the reaction-diffusion parameter 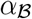 is varied. Below each complex, the flux enhancement achieved by the complex relative to delocalized catalysts is indicated, together with the average distance 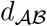 between 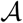 and 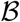 catalysts (in units of *r_c_*). For the 3D case, the conjectured optimal geometries matching the polyhedral solutions of the Thomson problem are shown (see main text).

We also sought to identify optimal 3D geometries. The increased configurational search space made the optimization slow and computationally challenging due to local optima. The best-performing geometries that we identified consisted of 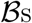 arranged on the surface of a sphere around 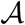, with radius that increases with 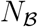 (see Fig. 5). In contrast to the 2D case, we never observed the separation of a second outer 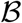 population in 3D, although we cannot rule out that this still occurs at higher 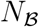. Except in special cases 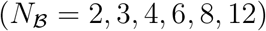 it is not possible to arrange points on the sphere such that all edges are of equal length; thus the 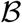 are not all equidistant from their neighbors. Interestingly, the optimal configurations generated by our numerical approach were similar to solutions of the well-known Thomson problem from classical electrostatics [52], where the objective is to minimize the electrostatic interaction energy of identical point charges on the surface of a sphere. Here, however, instead of interactions being defined by an identical local potential around each charge, the catalysts in our model effectively interact via the metabolite concentration fields, which depend on the positions of all catalysts. When we calculated the reaction flux for the exactly known configurations of the Thomson problem, we found that these always achieved a slightly higher reaction flux than any configuration found during our numerical optimization. Additionally, when we initialized the random search optimization with the solution of the Thomson problem, the algorithm was not able to identify any better configuration. We therefore conjecture that the Thomson problem configurations also optimize the reaction flux for the 3D model described here, provided that the radius of the sphere on which the 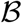 are arranged is chosen optimally.

Fig. 5 represents a minimal model, which illustrates basic physical principles governing optimal arrangements of catalytic particles. This model does not include an additional feature exhibited by many experimental systems: an intrinsic anisotropy of the catalysts. For instance, enzymes are reactive only at specific active sites rather than over their entire surface. With active sites, the relative orientations become additional degrees of freedom in catalyst arrangements. To explore the behavior of anisotropic catalysts, we introduced model catalysts with a reactive patch covering 1/6 of their surface (see Supplementary Information). We repeated the analysis of Fig. 3a for this model, to probe the effect of catalyst anisotropy on the average pathway flux obtained with different localization strategies. While we observed quantitative differences, the anisotropy did not qualitatively alter the relative performance of the localization strategies (Fig. S12). However, it did alter the symmetries of optimal catalyst complexes (Fig. S13). Despite the altered symmetries, the underlying design principle appears to be the same: balancing the advantageous effects of proximity and confinement against the detrimental effects of shielding and intermediate depletion.

## DISCUSSION

### Trade-offs

Our analysis of minimal models for spatially organized catalytic particles illustrated physical principles that are more general than the model assumptions: We found that two generic trade-offs govern the total reaction flux achieved by a given spatial arrangement. The first is fundamentally a trade-off between conversion efficiency and local depletion: Placing consecutive catalysts closely together, e.g. in larger clusters, increases the efficiency at which intermediate products are converted into final product, but excessive accumulation of catalysts depletes the primary substrate locally (Figs. 3a,b and S10). The second trade-off is between substrate shielding and intermediate confinement: Clustering of catalysts leads to shielding of interior catalysts (Fig. 4e), limiting their access to substrate (Fig. 4g). On the other hand, confinement of intermediates produced within the cluster (Fig. 4f) increases the number of potential interactions with downstream catalysts, and hence the apparent efficiency (Fig. 4h). The interplay between these trade-offs produces the phase diagram of optimal stoichiometries and spatial organization in Fig. 3e and the intricate optimal geometries of catalyst complexes in Fig. 5. While the latter depend also on the structure of the cataysts (Fig. S13), our qualitative findings are insensitive to these microscopic properties (Fig. S12). For the two fundamental trade-offs and all of our results the loss of intermediates is essential, but the precise loss mechanism does not qualitatively affect the behavior (Fig. S4).

### Design principles

The relative timescales of metabolite diffusion and catalytic reactions determine which side of each of the trade-offs should be favored in order to maximize the pathway flux, and therefore which type of localization strategy is preferable. In the reaction limited regime (slow reactions or fast diffusion), it is beneficial to form large clusters of catalysts, thereby favoring efficient transfer of intermediates between catalysts over access to substrate of the first catalyst in the pathway. This regime is probed by an *in vivo* experiment [53], in which three different types of RNA scaffolds were used to arrange the enzymes acyl-ACP reductase and aldehyde deformylating oxygenase into pairs, 1D strings, and 2D sheets. While the enzyme pairs only negligibly increased the production of the product pentadecane, the enzyme strings and sheets achieved an enhancement of ~80%. We simulated these experiments within our model (Fig. 6), finding a similar behavior of the reaction fluxes for the different arrangements in the reaction-limited regime. This also revealed that the sheet arrangement already behaves qualitatively similarly to the cluster strategy considered in Fig 3b, exhibiting the same signatures of the trade-off between substrate depletion and efficient intermediate transfer.

**FIG. 6:**
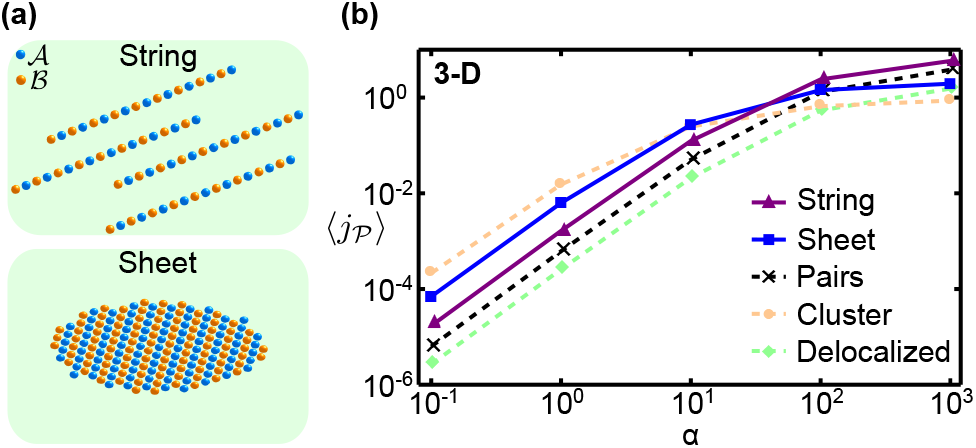
Analysis of the string and sheet localization strategies studied experimentally in [53]. (a) Illustration of the string and sheet configurations with alternating 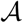 and 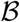 catalysts. (b) The pathway flux achieved by catalysts arranged into strings and sheets (solid lines) in comparison to paired, randomly clustered, and delocalized catalysts (dashed lines) for 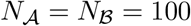.

In the diffusion limited regime (fast reactions or slow diffusion), clustering is detrimental. This regime applies to the experiment of Grotzky *et al.* [54], who found a decreased reaction flux when the diffusion-limited enzyme superoxide dismutase (SOD) was coclustered with HRP. Our model suggests that these catalysts should be arranged in pairs or small complexes, to ensure that the first catalyst receives a sufficient supply of substrate, while still achieving relatively efficient conversion of intermediate.

Several *in vitro* studies reported increases in the rate of product formation after positioning consecutive enzymes in close proximity [18, 19, 55, 56]. However, in such experiments the intermediate products are typically not subject to any loss to the surrounding environment or to competing reactions. In this setting, any enhancement effect will only be transient, whereas steady-state fluxes will not be affected by the spatial organization. Indeed, no enhancement was observed when the sequential enzymes GOx and HRP were fused together using a small molecular linker [20]. The authors attributed the earlier reports of flux enhancement [19] to local changes in pH around DNA-based scaffolds that increase the enzymatic activities of GOx and HRP. Furthermore, no enhancement was observed when catalase was included as a scavenging enzyme that consumes the intermediate H_2_O_2_. For realistic parameters and concentrations of GOx, HRP, and the scavenging enzyme as used in [20], our theoretical framework also predicts a negligible enhancement in reaction flux (Fig. S7), consistent with these observations. To achieve a significant flux enhancement via proximity alone would require either a strongly reduced concentration of the GOx-HRP enzyme pairs, or a significantly increased concentration of the scavenging enzyme (see Supplementary Information).

### Steric effects

The second trade-off is the result of steric effects arising from the discrete nature of catalysts. Our model illustrates that these steric effects can have a strong impact, altering reaction fluxes up to 4-fold. With our sperical catalysts, the steric effects are smaller in 3D than in 2D, largely due to the smaller maximal packing density in 3D. The effect of catalytically inert crowding agents on the reaction fluxes can be predicted based on our results for reactive catalysts: The effect of crowding agents on the first reaction flux is similar to that of the second catalysts, since both shield the substrates from accessing the first catalysts. In contrast, the effect of crowding agents on the second reaction should to be weaker, since the confinement of intermediates is only advantageous if the number of collisions between intermediates and the second catalyst is increased. With inert particles as crowding agents instead of the second catalysts, this collision number is increased less significantly.

Furthermore, in enzymatic systems the catalysts are reactive only at a specific active site rather than over their entire surface. Introducing such an active site adds the orientation of the catalysts as additional degrees of freedom to localization strategies. However, we found that orientation effects were less important than proximity in determining the flux generated by different strategies (Fig. S12). We also found that anisotropic catalyst reactivity breaks the symmetry of optimal arrangements within a model cluster (Fig. S13). However, the effects of substrate shielding and intermediate confinement still appear to play a vital role in shaping the optimal configurations leading to qualitatively similar arrangements in which one fraction of the 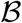 catalysts are placed close to a central 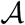 with the other fraction further away. Thus the organization principles outlined here are largely robust to the microscopic details of catalysts.

### Generalized Thomson problem

We saw that the interplay of two fundamental tradeoffs leads to complicated and varied optimal geometries for model multi-catalyst complexes with a single catalyst of the first type surrounded by several catalysts of the second type. Notably, the resulting optimal configurations show striking similarities to the well known Thomson problem of classical electrostatics. However, in contrast to the Thomson problem, where electrons interact via the Coulomb potential and the total energy function is determined as a sum of the individual pair interactions, for our model of multi-catalyst complexes the metabolite concentration fields mediate effective many-body interactions between catalysts. Consequently, the total reaction flux cannot be expressed simply as a superposition of the individual contributions of the catalysts. Therefore, the problem of finding the optimal catalyst complex arrangements adds an additional level of complexity to the class of generalized Thomson problems. Although these problems are very easy to pose they are notoriously difficult to solve rigorously. For the standard Thomson problem the symmetries have only be rigorously identified for a few electron numbers [57–59] and solving the general case has been included in the list of eighteen unsolved mathematical problems for the 21st century by Steven Smale [60]. Besides the interesting mathematical nature of the stated problem and the symmetries that emerge, the optimal structure of the model complex may also provide valuable insights into the design principles of such complexes. Indeed, similar optimization problems, yet with very different objectives, were previously identified as useful minimal models for understanding the structure and geometry of biological materials ranging from proteins and viral capsids to plant phyllotaxis and honeycombs in beehives [1, 61–66].

## METHODS

### Reaction-diffusion model

We modeled the metabolites 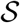 and 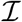 as small molecules that move by diffusion. Their concentrations, 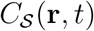 and 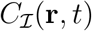 respectively, follow

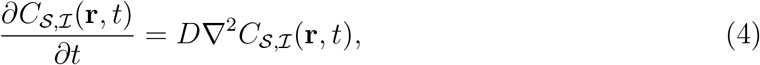

where *D* is the diffusion coefficient, which we assumed to be the same for both metabolites. Since the catalysts are mesoscopic objects (macromolecules, organelles, or cells), we describe them as discrete spatially-extended reaction centers (see Fig 1a). We implemented the reactions of Eq. 1 through boundary conditions imposed on the metabolite concentrations at the surface of the respective catalyst, 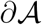 or 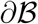,

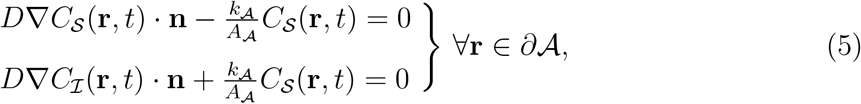

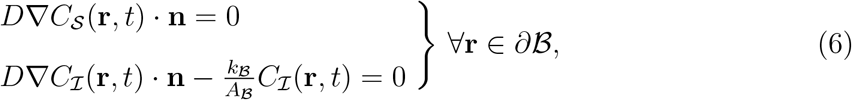

where 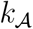 and 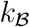 are the intrinsic catalytic efficiencies of 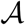 and 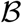 respectively, 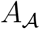 and 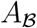 the catalyst surface areas, and **n** is a unit vector normal to the surface. Eq. 5 represents the conversion of 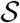 to 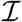 catalyzed by 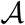. Eq. 6 describes the consumption of 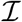 by 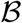, as well as a no flux condition for 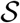 at the surface of 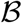, since there is no reactive interaction between 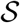 and 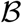. In Eqs. 5, 6 we have neglected saturation of the catalysts, assuming we are in the low metabolite concentration regime. We study the relaxation of this assumption in the Supplementary Information.

The intrinsic catalytic efficiencies 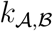 in Eqs. 5, 6 are effective parameters that describe reactions for metabolites in contact with a catalyst. For systems of collaborative microorganisms or organelles, these parameters will depend on the metabolite uptake rate through transporter protein concentrations and activities and the membrane permeability, as well as the turnover rate of metabolites within the cell or organelle. For enzymatic reactions the *k*’s will instead be determined by microscopic details such as the interaction potential between metabolite and catalyst, and the transition path between the substrate-catalyst to product-catalyst complexes. Importantly, these parameters differ from the macroscopic catalytic efficiencies that would be measured in solution, which take into account not only the activities of the catalysts, but also the timescale of metabolite-catalyst encounters via diffusion. The correspondence between the macroscopic catalytic efficiency, *k*, and the intrinsic efficiency, *k*, is usually modeled as 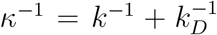 [67, 68], where *k_D_* is the rate at which substrates arrive at a catalyst via diffusion. Reactions can broadly be classified as reaction-limited when *k* < *k_D_*, in which case the macroscopic efficiency is *κ* ≃ *k*; or diffusion-limited when *k* > *k_D_*, in which case *κ* ≃ *k_D_*. The macroscopic catalytic efficiency therefore has an upper bound determined by the diffusive arrival rate, which for enzymatic reactions is *κ* ≾ *k_D_* ≈ 10^9^ M^-1^ s^-1^. It has been shown that the intrinsic association rate of the enzyme-substrate complex, which determines the intrinsic catalytic efficiency in the fast reaction regime, can reach *k* ~ 10*k_D_* [69], although this value may be exceeded depending on the specific interaction potential and effective interaction radius.

The reaction cascade is supplied with substrate from the surrounding environment. We assume the environment to provide a homeostatic level *C*_0_ of 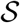, whereas the environmental concentration of intermediates remains negligible, corresponding to the conditions

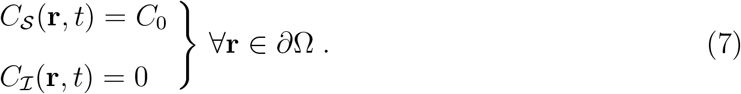

at the system boundary, *∂*Ω. As a measure of the collective performance of the catalysts we focused on the steady-state production rate of 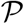,

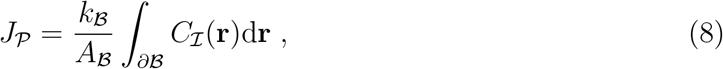

where the integral is taken over the surface of all 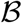 catalysts. Analogously, we define the rate of 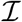 production as,

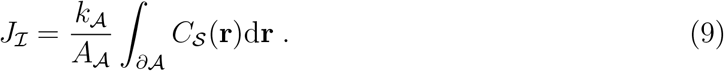

In this work we are primarily interested in how the spatial organization of sequential catalysts influences the pathway flux 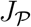. In general, however, 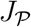 will also depend on the shape and size of the catalysts and the system geometry *δ*Ω, in addition to all the model parameters. For simplicity we assumed that 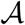 and 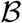 have the same size and a spherical shape, such that 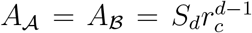 in a model of dimension *d* with *S_d_* the surface area of a d-dimensional unit sphere. We also assumed a spherical system geometry with a fixed radius 100 *r_c_* for the system boundary. Note that taking *r_c_* = 2nm on the typical scale of a single enzyme molecule results in an effective enzyme concentration in the higher nanomolar range for the small values of 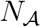 and 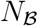 considered here, which is within the range expected under intracelullar conditions. Rescaling all lengths with the interaction radius, *r_c_*, we identify from Eqs. 5, 6 two dimensionless reaction-diffusion parameters, 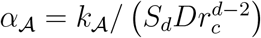 and 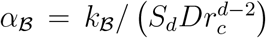, that together with the dimensionless system radius determine the metabolite concentration profiles. Finally, since Eqs. 4–6 are linear in the metabolite concentrations, these can be normalized by 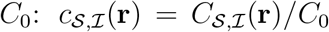. Rewriting 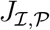 in terms of these dimensionless variables, we identify the dimensionless reaction fluxes

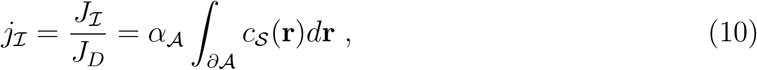

where 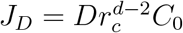, and 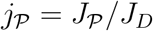 as given by Eq. 3.

Fluxes for different catalyst arrangements were calculated by numerically solving the steady-state nondimensionalized versions of Eqs. 4–7 using COMSOL Multiphysics (COMSOL AB).

### Catalyst arrangement ensembles

The distributions of pathway fluxes for different model parameters and localization strategies were determined in each case by sampling an ensemble of 3000 random catalyst configurations and computing for each configuration the steady-state flux.

For the delocalized scenario, these configurations were generated by distributing the catalysts uniformly over the system. For a two dimensional spherical symmetric system a uniform distribution is achieved by picking for the center of each catalyst a radial position 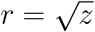 where *z* is uniformly distributed over the interval 0 ≤ *z* < (*R* — *r_c_*)^2^, and an angular position *θ* from the interval 0 ≤ *θ* < 2*π*. Similarly, in three dimensions the radial position is *r* = *z*^1/3^, and the angular coordinates 0 ≤ *θ* < 2*π* and *ϕ* = arccos(2*v* — 1) where 0 ≤ *v* ≤ 1. After distributing all catalysts in this way, we tested whether any two catalysts overlapped. If any pair had a distance smaller than 2*r_c_* between their centers, we moved this pair away from each other, along the line connecting the centers, until their separation was larger than 2*r_c_*. After all overlapping pairs were relocated, the procedure was repeated to avoid overlaps created by the repositioning.

For the different studied catalyst organizations, we also considered ensembles of configurations generated by a similar procedure. In the case of 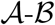 pairs and the complex configurations in Fig. 3e, their centers were distributed randomly over the system as described above. The center-center distance between catalysts within a pair or complex was fixed at 3*r_c_*, while their orientation was chosen randomly. In the case of clustered catalysts, a center position for the cluster within the system was chosen randomly. The catalysts were then randomly positioned within a circular (in 2d) or spherical (in 3d) region so as to achieve a packing density of *ϕ* = 60% in 2D or *ϕ* = 50% in 3D. In all cases, cycles of rearrangements were made in order to avoid catalyst overlaps.

### Cluster arrangement optimization

To determine the optimal 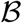 configuration around a single 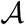 localized at the system center, we used a Monte Carlo optimization algorithm to iteratively explore the catalyst configuration space.

The optimization algorithm was initialized with a random configuration of 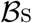. From this configuration a new trial configuration was sampled by selecting one 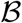 at random and moving it a distance *l* in a random direction to a new position. If this trial configuration led to an increase in reaction flux 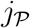 it was accepted and used as the starting configuration for the next trial step, otherwise it was rejected and a new trial was generated from the previous best configuration. This procedure was repeated until a termination criterion of either a defined total number of iterations (set to 10^4^), or a number of successive non-improving iterations (300–400), was reached. We made two further modifications to this basic algorithm that were found to speed up convergence of the optimization. First, after a trial step was accepted the subsequent trial step was taken in the same direction. Second, the step length was decreased during the course of the optimization process.

In general, this procedure does not guarantee a convergence to the global optimum. We therefore performed 30 realizations of the optimization procedure, with different initial configurations, for each set of model parameters. Most realizations resulted in the same final configuration, which we are confident to be the global optimum.

## Acknowledgments

The authors thank Erwin Frey, David Nelson, and Aleksandra Walczak for useful discussions, and Bernhard Altaner and Giovanni Giunta for comments on the manuscript. This work was supported by the German Excellence Initiative via the program “Nanosystems Initiative Munich” and the German Research Foundation via SFB1032 “Nanoagents for Spatiotemporal Control of Molecular and Cellular Reactions”. F.H. was supported by a DFG Fellowship through the Graduate School of Quantitative Biosciences Munich (QBM).

## Author contributions

All authors designed the research. F.H. performed the research. F.H., F.T. and U.G. analyzed the results and wrote the paper.

## Supplementary Information for

### I. CORRELATIONS OF THE REACTION FLUX WITH DIFFERENT PROPERTIES OF THE CATALYST ARRANGEMENTS

In Fig. 2 of the main text we saw that the reaction flux 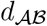 is correlated with the mean distance 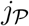 between the catalysts 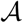 and 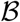 in random delocalized arrangements. For small values of the reaction-diffusion parameter *α*, the reaction flux 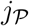 is strongly anti-correlated with the distance 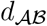, while we found a somewhat weaker but positive correlation in the large a regime. Figure S1 shows that similar correlations exist between 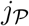 and the mean distance between catalysts of the same type (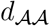 and 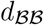) and between 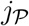 and the mean radial coordinates of catalysts (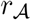 and 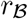).

**FIG. S1:**
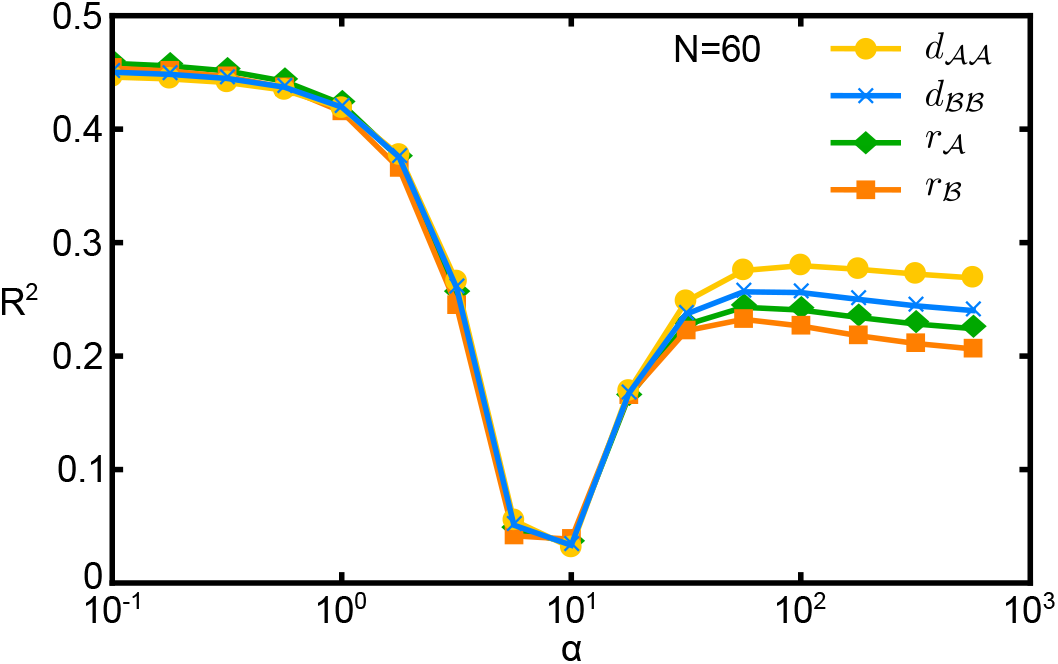
Coefficient of determination (*R*^2^) of the linear regression of the reaction flux 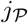 against different geometrical characteristics of random delocalized catalyst arrangements: the mean distance 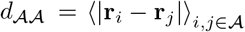 between 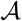 catalysts, the mean distance 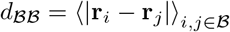 between 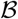 catalysts, the mean distance 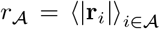 of 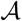 catalysts from the center, and the mean distance 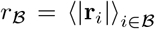 of 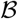 catalysts from the center. In each case the degree of the correlation (*R*^2^) displays a similar dependence on the reaction-diffusion parameter *α* as observed in Fig. 2c of the main text for the mean distance 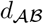 between the catalysts 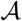 and 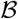.

### II. SLOW LEAKAGE OF INTERMEDIATES

For the examples shown in the main text figures, we assumed that the intermediates are lost as soon as they reach the system boundary (absorbing boundary condition, 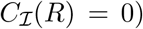. This assumption of fast leakage can be relaxed by introducing a boundary condition that is partially absorbing and partially reflecting,

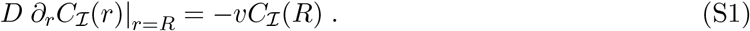

Here, *v* quantifies the permeability of the boundary to intermediates. By dividing Eq. S1 by *D* and rescaling all lengths by *r_c_* we obtain the non-dimensional leakage parameter

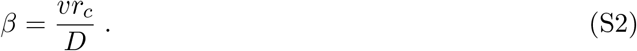

For very slow leakage (*β* ≪ 1), almost all intermediates are converted into product and the overall pathway flux is essentially equal to the production rate of intermediates, 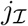. The latter still depends on the catalyst arrangement, because the net flux of 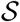 into the system varies with the rate of the first reaction, which depends on the extent of substrate depletion. Thus catalyst pairs and delocalized catalysts perform almost identically, since depletion is negligible in these cases (Fig. S2a). Clusters lead to stronger depletion and therefore still display a reduced flux. For larger 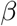, on the other hand, it is vital that intermediates are rapidly transferred between the catalysts to prevent their loss. For instance, at 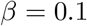, the system behavior (Fig. S2c) is already similar to the absorbing boundary limit (Fig. 3a of the main text).

We observe that the crossover between the pathway fluxes of clusters and pairs shifts towards smaller *α* values as *β* is decreased (Fig. S2). This behavior can be understood as follows: For small leakage rates, intermediates can explore a larger fraction of the system and thus come in contact with a larger number of 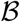 catalysts before leaking out. This reduces the beneficial effect of catalyst proximity and since the detrimental effect of substrate depletion remains unchanged, the cluster strategy starts to be outperformed by the pair strategy already at smaller *α* values.

In the main text we saw that the enhancement of the catalyst pair strategy compared to delocalized catalysts increases as the number of such pairs in the system is reduced (Fig. 3c and d). This trend persists for different leakage rates, but overall, the enhancement decreases when the leakage rate is decreased (Fig. S3). As discussed in the previous section, for small leakage rates intermediates that are not rapidly converted by the closest 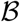 catalyst have a higher probability to react with any other 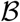 catalyst in the system, which reduces the beneficial effect of proximity, and hence the flux enhancement.

**FIG. S2:**
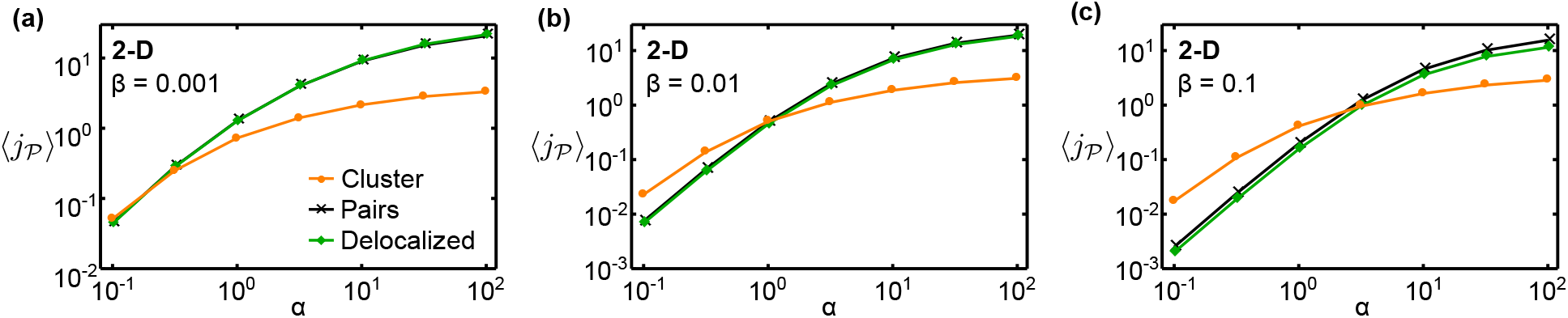
Influence of intermediate leakage on the pathway flux. Comparison of the fluxes for delocalized, paired, and clustered catalysts for three different leakage parameters, (a) *β* = 0.001, (b) *β* = 0.01, and (c) *β* = 0.1. Besides the leakage parameter, all other aspects of the model are chosen as for Fig. 3a of the main text.

**FIG. S3:**
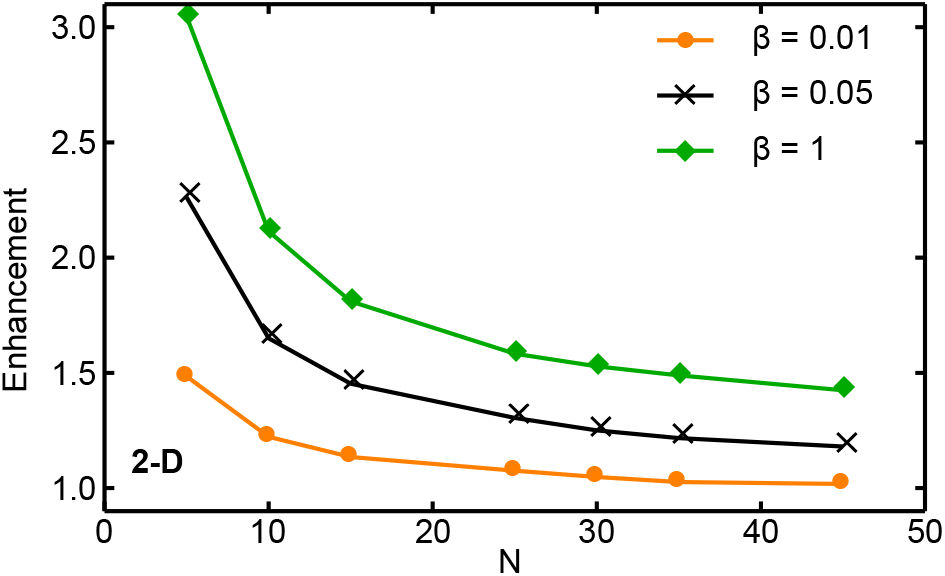
Flux enhancement achieved by catalyst pairs relative to delocalized catalysts. The enhancement is plotted as a function of the number of catalysts, 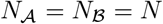, for three different values of the leakage parameter *β*. All other aspects of the model are chosen as for Fig. 3c of the main text.

### III. SUBSTRATE PRODUCTION AND INTERMEDIATE LOSS WITHIN THE SYSTEM

For the examples shown in the main text figures (and the previous sections of this Supplementary Information), we assumed that the influx of substrates and the loss of intermediates takes place at the system boundary. In this scenario, the reaction pathway is supplied with substrate 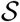 from the surrounding environment, in which a constant concentration *C*_0_ of 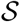 is maintained. The intermediates 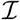, in turn, are lost to the surrounding environment where their concentration is negligible. As an alternative we will consider here a scenario in which these processes take place within the system: 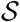 is maintained at a homeostatic level inside the system, while 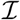 is lost either due to an intrinsic instability of the molecule or due to competing reactions. In this section, we study how the fluxes achieved by the different localization strategies behave for the alternative scenario and whether the choice of scenario affects the qualitative behavior of the model.

To determine the fluxes for the alternative scenario, we include reaction terms for the production of 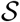 and the decay of 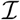 into the diffusion equations (Eq. 4) of the main text. This leads to the steady state conditions

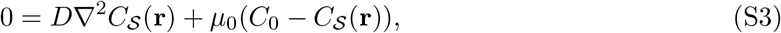

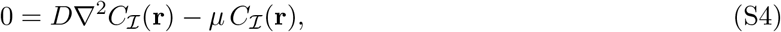

where *μ*_0_ is the rate at which 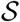 relaxes to the homeostatic level *C*_0_, and *μ* is the rate at which spontaneous decay or competing reactions consume 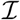. At the system boundary we now impose no-flux boundary conditions for both 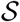 and 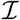. By dividing Eq. S4 by *D*, and by rescaling all lengths by *r_c_*, we obtain the loss parameter

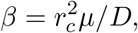

which measures the strength of the loss mechanism. This dimensionless parameter corresponds to the square of the ratio of the catalyst radius, *r_c_*, to the distance 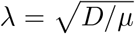 that an intermediate typically diffuses before it is lost. Equivalently, it can be interpreted as the ratio of two timescales, a typical diffusion time over a distance ~ *r_c_* and a typical decay time, 1/*μ*, for the loss mechanism. Note that this parameter differs from the leakage parameter *β* defined in the previous section, which measures the leakage of intermediates at the system boundary. Similarly, we obtain from Eq. S3 a second dimensionless parameter,

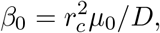

associated with the mechanism that supplies 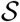, with analogous interpretations as the parameter *β*. For experimental systems, *β*_0_ will typically take on small values, since the diffusion timescale is much shorter than the timescale for the production of 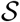 (which determines the value of *μ*). For our analysis, we therefore set the parameter *β*_0_ to a constant small value of 0.001 and consider different loss parameters *β* that are either smaller than, equal to, or larger than *β*_0_. Furthermore, we normalize the pathway flux by V*μ*_0_*C*_0_, such that it becomes independent of the concentration *C*_0_.

Fig. S4 shows the pathway fluxes of the different localization strategies (delocalized, clustered, and pairs) as a function of the reaction-diffusion parameter *α* for different loss parameter values *β*, in both two and three dimensions. We find that the behavior of the fluxes is similar to the case of slow leakage of intermediates examined in the previous section. In particular, as *β* is decreased, the point at which the crossover between the cluster and pair strategies occurs is shifted to smaller *α* values, and the enhancement achieved by the pair strategy over the delocalized arrangement is diminished. This demonstrates that the fluxes behave qualitatively in the same way, irrespective of whether the processes of 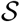 supply and 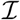 loss occur within the bulk or at the boundary of the system.

**FIG. S4:**
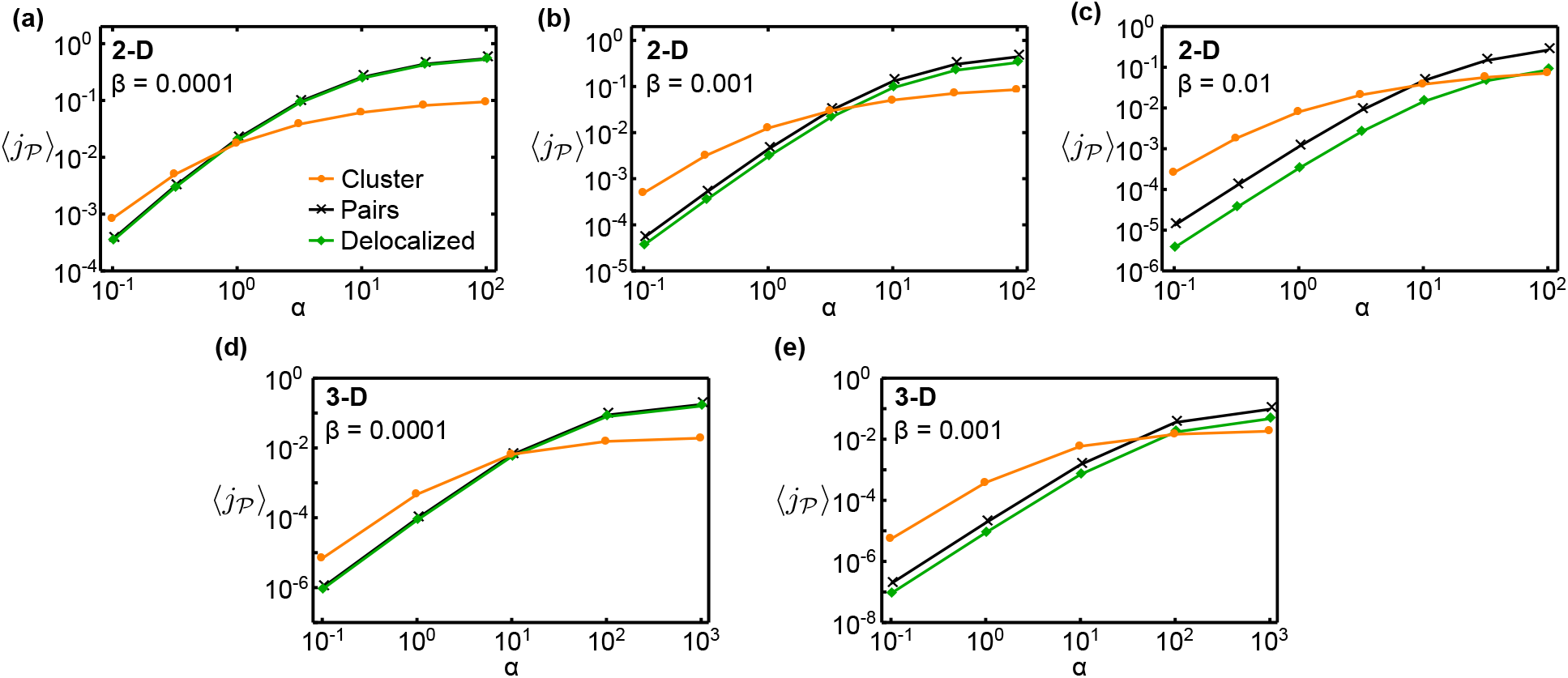
Comparison of the pathway fluxes for delocalized, paired, and clustered catalysts for the scenario where substrate production and intermediate leakage occur inside the system. The five panels show the mean pathway fluxes as a function of the reaction-diffusion parameter *α*, for several different values of the loss parameter *β* and in both two and three dimensions, as indicated. Panels (a), (b), and (c) are analogous to Fig. 3a in the main text, while panels (d) and (e) are analogous to Fig. 3b.

### IV. ANALYTIC APPROXIMATION OF THE ENHANCEMENT ACHIEVED BY THE PAIR STRATEGY

Following the theoretical approach of Fu *et al* [1], Wheeldon *et al.* [2] predicted that catalyst proximity has an advantageous effect on the reaction flux only when the production rate of intermediates by the first catalyst, *k*_cat_, is fast compared to the diffusivity, *D*, of the intermediate. This analysis is based on the fundamental solution of the diffusion equation that assumes diffusion over an infinite domain with an initial influx pulse of intermediates at time and position zero, 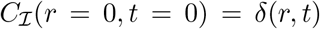, corresponding to a local intermediate production event by the first catalyst. Since the first catalyst constantly produces intermediates at discrete time points with a mean rate *k*_cat_ = 1/*τ*, the intermediate concentration profile was assumed to be the sum over the fundamental solution with the production time points shifted by *r*,

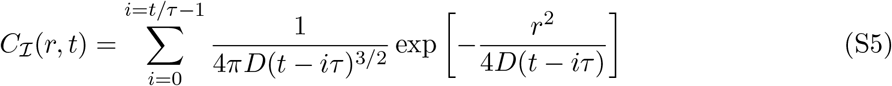

In steady-state, this results in a Gaussian concentration profile around the position of the first catalyst where the intermediates are produced. The ratio between *k*_cat_ and *D* determines how pronounced the maximum of the intermediate profile is around the first catalyst. This suggests that placing the second catalyst close to the first catalyst only has an effect on the reaction flux when the concentration peak around the first catalyst is significant, which is only the case for *D/k*_cat_, <1 μm^2^. However, this argument is based on assumptions that limit its applicability.

The assumption of a single source of intermediates in an infinite space is not applicable for systems with many catalysts in a finite space. The intermediates produced by one of the 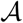 catalysts can react with any of the other 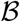 catalysts in the system, rather than just the closest 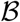. Furthermore, in a finite domain without any loss mechanism, intermediates will accumulate in the system to a level where the rate of product formation by the second catalyst equals the rate of intermediate formation by the first catalyst. Hence, formation of catalyst pairs will not enhance the steady state flux. The pathway flux can then only be transiently enhanced, in the time window before steady state is reached, during which intermediates have not yet accumulated to a constant level. Thus, it is essential to also explicitly include mechanisms for the loss of intermediates in the model scenario. It is worth noting that this applies also to experimental scenarios: In experimental studies where no loss mechanism is present, any observed localization-induced enhancement in the flux must either be transient, due to there being no intermediate products in the system initially, or be the result of altered catalyst activity caused by the localization procedure.

To obtain a theoretical understanding of the enhancement achieved by the pair strategy, we therefore derived an analytical expression for the enhancement of a catalyst pair in the presence of loss. To this end we considered a spherical domain of radius *R* containing a single catalyst pair, representative for one of the many catalyst pairs and the surrounding space within a larger system. The size of the domain is determined by the concentration *ρ* of catalyst pairs in the system via 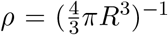. Inside the domain, the catalyst pair is positioned such that the 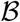 catalyst is localized at the center and the 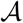 catalyst a distance *d* away from the center. The 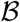 catalyst is treated as a spherical reaction center with radius *r_c_*, which is reactive over the entire surface. For the 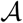 catalyst we exploit the spherical symmetry and assume that the production of intermediates occurs uniformly on a spherical shell of radius *d* around 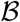. At the domain boundary at radius *R* we apply a reflective boundary condition on the intermediate concentration profile, which represents the exchange of intermediate with equivalent domains around other catalyst pairs. In the previous section we found that the qualitatve behavior of the model does not depend on the mechanism of intermediate loss. For analytic simplicity, we assume here that intermediates are lost everywhere in the system with a rate *μ,* rather than at the system boundary. The corresponding reaction-diffusion equation governing the dynamics of the intermediate is then

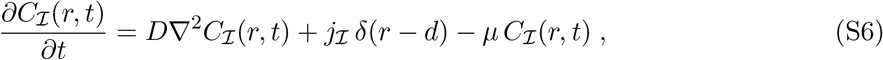

with the boundary conditions

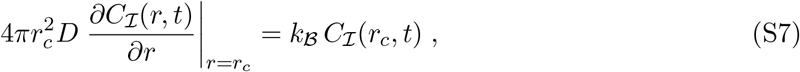

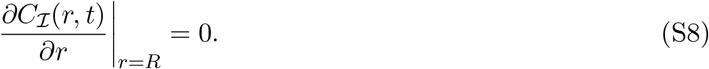

Here, 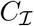 is the density profile of the intermediate 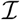, *δ* is the Dirac delta function, 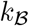 is the intrinsic reaction rate at the surface of 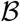, and *D* is the diffusion coefficient of 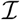. In steady-state we can solve this reaction-diffusion system analytically and determine the reaction flux of a single catalyst pair,

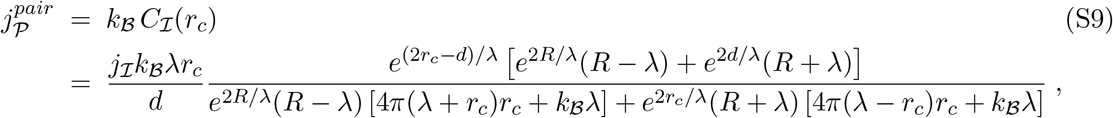

where λ = (*D/μ*)^1/2^ is the leakage length scale, which measures the distance a metabolite typically diffuses before it is lost.

To compute the enhancement achieved by the pair strategy, we have to relate the reaction flux of a catalyst pair in proximity, 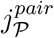, to the reaction flux of a delocalized catalyst pair, 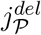.

**FIG. S5:**
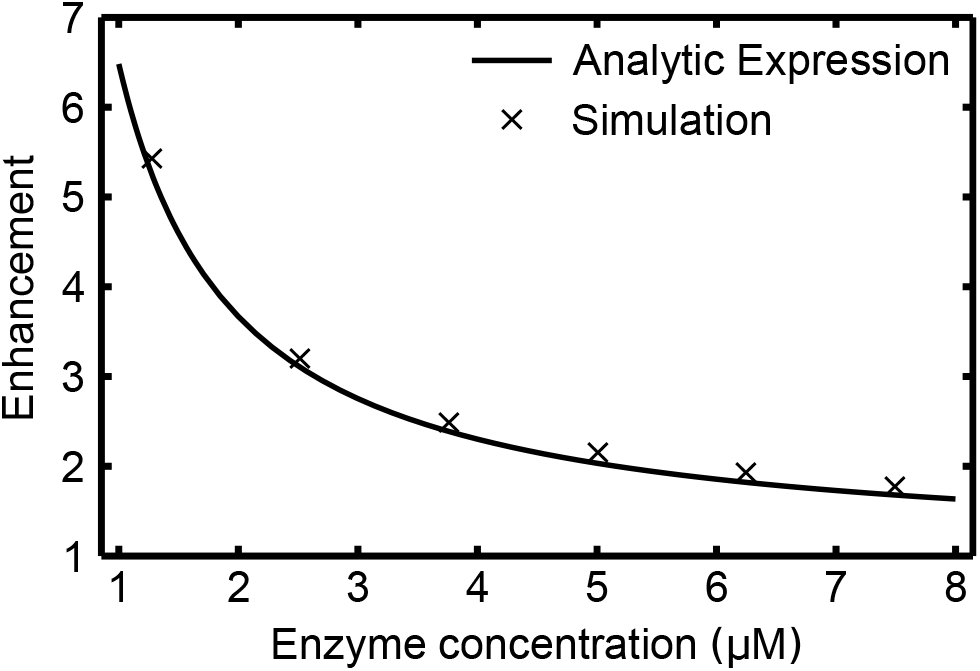
Comparison between the analytical approximation of the flux enhancement for the pair strategy (leakage length λ = 0.1 μm, distance between enzymes *d* = 2nm) and the corresponding numerical computation based on the full model with the corresponding parameters (*r_c_* = 2 nm).

The latter flux can be approximated by assuming that the distance *d* between the two catalysts in the pair corresponds to the typical distance, *R*, between catalysts in the delocalized scenario, 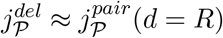. With this we obtain the enhancement factor

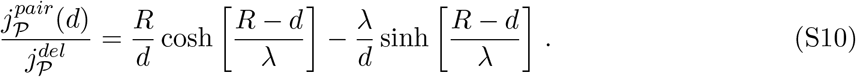

Fig. S5 shows that this analytical approximation agrees well with the numerically computed enhancement factor of the full model with intermediate loss within the system. The enhancement (S10) depends only on the distance *d* between paired catalysts, the concentration of catalyst pairs via 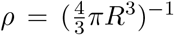, and the leakage length scale λ, but is independent of the reaction rate 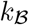. To understand why this enhancement is independent of 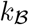, consider that the reaction probability, 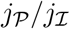, is composed of two factors. The first is the probability that an 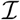 molecule produced by 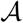 reaches 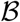 by diffusion, which will depend on the positioning of the catalysts relative to one another. The second is the probability that an 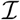 molecule is processed to 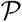 given it reached 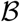. Only the latter probability depends on the reaction rate of 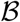, and since this probability is the same for delocalized catalysts and catalyst pairs, the enhancement becomes largely independent of the reaction rate.

**FIG. S6:**
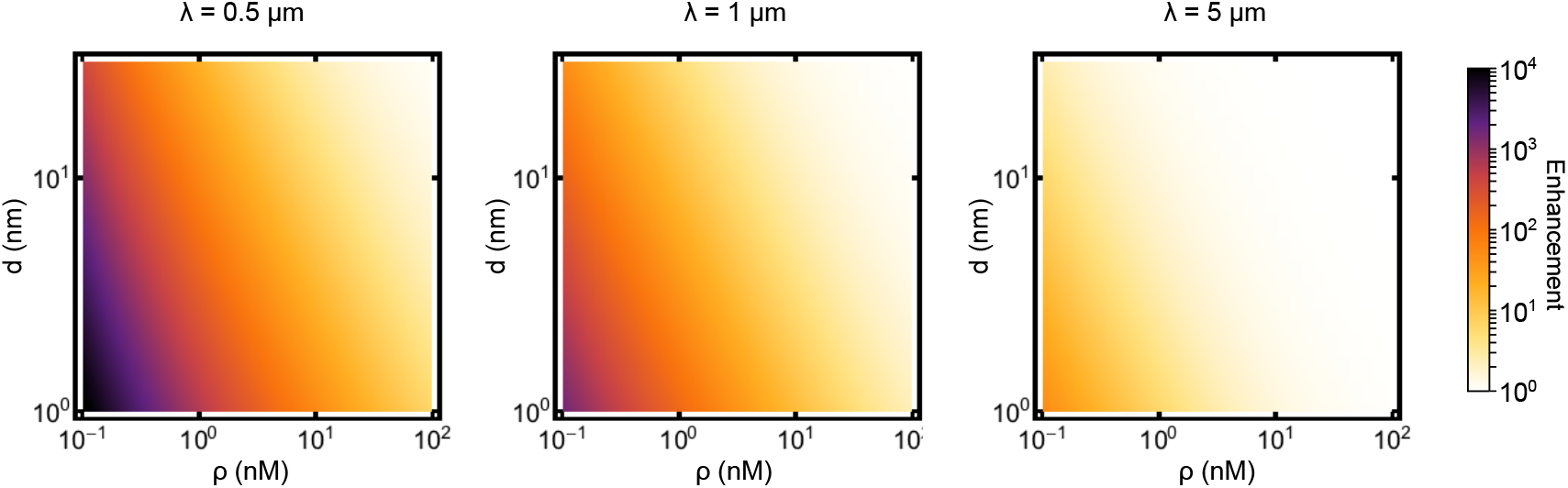
Analytic approximation of the flux enhancement achieved by the pair strategy. The enhancement is plotted as a function of the catalyst pair concentration, *ρ*, and the distance between the catalysts, *d*, for three different leakage lengths λ.

We can see from Eq. S10 that the largest enhancement can be achieved for small inter-catalyst distances *d*, small catalyst pair concentrations *ρ* and small leakage length scales λ (see Fig. S6). The analytical expression (S10) is also useful for the interpretation of the results of experimental studies. To illustrate this, we discuss the experimental findings of Zhang *et al.* [3] who conjugated the consecutive enzymes glucose oxidase (GOx) and horseradish peroxidase (HRP). The conjugation was performed with a molecular linker that was shown not to alter the intrinsic activity of the enzymes. The enzyme catalase was used as a scavenging enzyme that competes for the intermediate HO of the GOx-HRP pathway. The concentration of catalase, [*CAT*], then determines the leakage length via 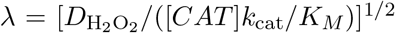, where *k*_cat_/*K_M_* is the catalytic efficiency of catalase, and *D*_H_2_O_2__ is the diffusion coefficient of H_2_O_2_ In the experiment by Zhang *et al.,* no enhancement could be detected for an enzyme pair concentration of *ρ* ~1nM, an interenzyme distance of *d* ~2nm, and three different catalase concentrations (380 nM, 190 nM, and 19 nM). Eq. S10 predicts enhancements of 16% for 380 nM catalase, 8% for 190 nM catalase, and 0.8% for 19nM catalase (see Fig. S7), which are indeed rather small and hardly detectable experimentally. To achieve a significant enhancement (e.g., larger than two fold) one would need to either reduce the concentration of the enzyme pair or increase the concentration of the competing enzyme (catalase), as indicated in Fig. S7.

**FIG. S7:**
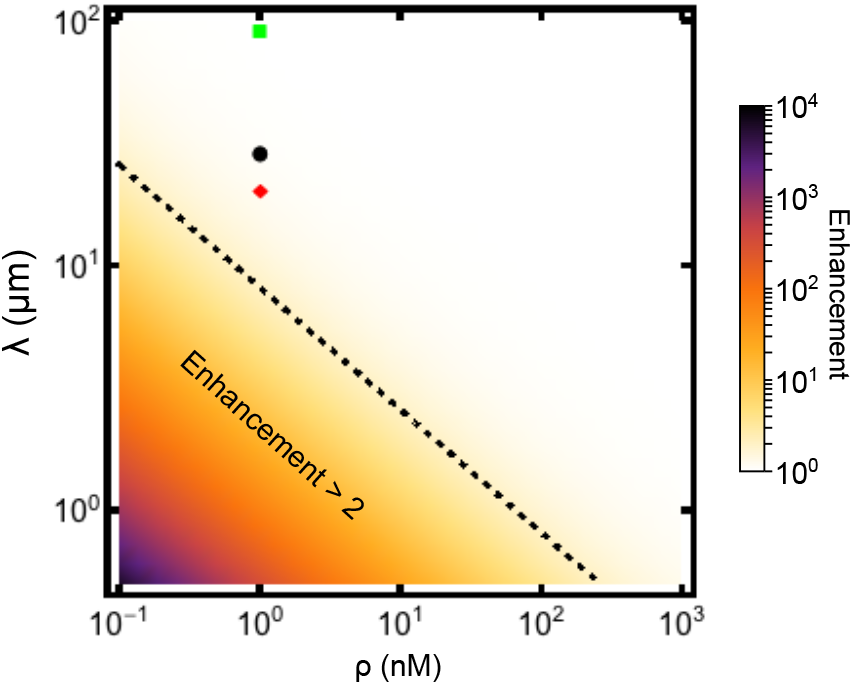
The experimental conditions of Zhang *et al.* [3] marked in a color map plot of the analytically predicted flux enhancement as a function of the leakage length λ and the enzyme pair concentration *ρ*, with a fixed interenzyme distance of *d* =2 nm. The three markers show the conditions corresponding to the three different catalase concentrations used in [3], [*CAT*] =19nM (green square), [*CAT*] =190nM (black circle), and [*CAT*] =380nM (red diamond). Together with the catalytic efficiency of catalase, *k_cat_/K_M_* = 6.4 x 10^6^ M^-1^ s^-1^, and the diffusion coefficient of H_2_O_2_, *D* = 1000 μm^2^ s^-1^, we determined the leakage length via λ = [*D*/([*CAT*]*k_cat_/K_M_*)]^1/2^.

### V. NONLINEAR REACTION KINETICS: SATURATION EFFECTS

#### Nonlinear model

In the main text we assumed that the catalytic reactions are linear in the metabolite concentrations 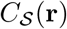 and 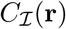. However, in general such reactions follow a nonlinear Michaelis-Menten-like scheme, in which the reaction becomes saturated for high metabolite concentrations. To study the impact of catalyst saturation, we extended our model by replacing the reaction boundary conditions, Eqs. (4) and (5) in the main text, with

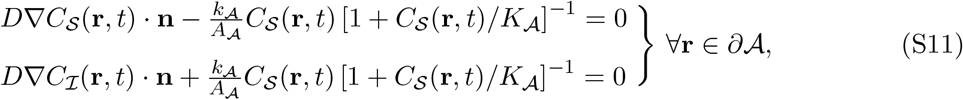

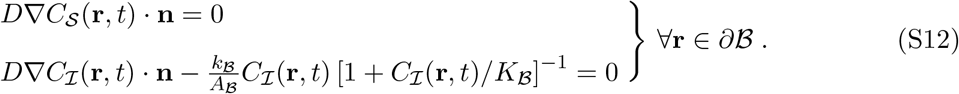

After rescaling all metabolite concentrations by the substrate concentration *C*_0_ in the environment, we obtain two additional non-dimensional parameters, measuring the degree of saturation of the two catalytic reactions, 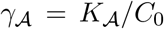 and 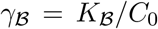. When both 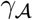 and 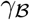 are large, we recover the limit of linear reactions studied in the main text.

#### Behavior for symmetric saturation

We first consider the case where both reactions have the same value for the saturation parameter, 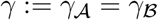, and analyze the behavior of the model as *γ* is decreased, corresponding to more saturated catalysts. For *γ* = 1, where saturation effects are expected to set in, Fig. S8a shows the mean reaction flux 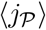 as a function of the reaction-diffusion parameter *α*. As in Fig. 3a of the main text, Fig. S8 compares the clustered arrangements with the 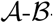-pair arrangements and the delocalized arrangements. Note that Fig. S8 and Fig. 3a differ only in the boundary conditions used to compute the respective data, i.e. Eqs. (S11) and (S12) instead of Eqs. (4) and (5) of the main text. With *γ* = 1, the behavior remains almost indistinguishable from that in Fig. 3a. However, with a tenfold lower saturation parameter (*γ* = 0.1), the crossover between the cluster and pair strategy is noticeably shifted towards larger *α* values (Fig. S8b). Nevertheless, the qualitative behavior remains the same as for the linear reactions in Fig. 3a of the main text.

This behavior can be understood by analyzing the impact of catalyst saturation on the competing effects of substrate depletion and efficient intermediate transfer. The nonlinear kinetics mitigates the effect of substrate depletion in two ways. First, when catalysts are saturated the fraction of substrate molecules in their vicinity that are consumed is reduced, which reduces the local depletion around the 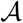 catalysts. Second, even when the substrate is locally depleted, this produces significant changes in the reaction flux only if the concentration is reduced below the saturation threshold of the catalysts. Thus, as 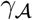 is decreased, the point at which substrate depletion has a significant effect on the flux is shifted to larger a values.

**FIG. S8:**
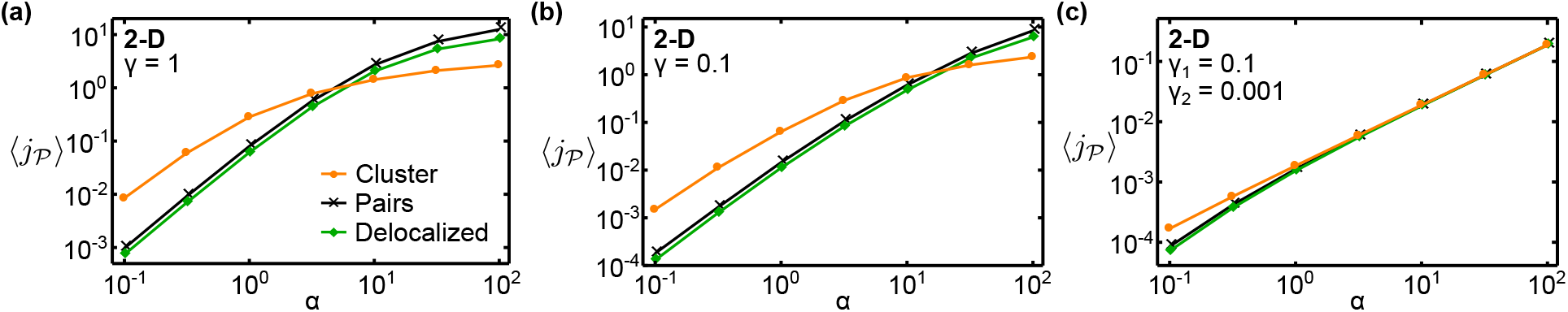
Effect of catalyst saturation on reaction fluxes for different catalyst arrangements. As in Fig. 3a of the main text, we compare the mean reaction fluxes 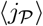 produced by three different spatial strategies of arranging catalysts: clustered (orange), 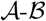-pairs (black), and delocalized (green), in each case with the catalyst copy numbers set to 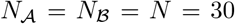. The fluxes are plotted as a function of the reactiondiffusion parameter 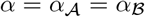. (a) 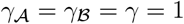, (b) 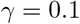, (c) 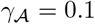 and 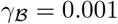.

In contrast, for the second reaction, intermediates leak out of the system, which prevents them from accumulating to a level where the 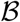 catalysts become saturated. Hence, the reaction kinetics of 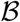 catalysts stays approximately linear, such that the 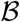 catalysts are able to exploit the increased local intermediate concentration that is conferred by proximity. All these effects tend to reduce the downsides and amplify the benefits of clustering catalysts, thereby extending the regime where clustering is beneficial to larger *α* values. For small 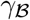, however, where the 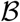 catalysts are completely saturated, the catalyst arrangement has essentially no effect on the pathway flux (Fig. S8c). In this case all localization strategies achieve the same flux, except for small a, where clustering is still advantageous.

The above picture applies in the regime where the loss of intermediates is relatively rapid. If intermediate leakage is reduced, the beneficial effect of placing 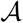 and 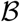 catalysts closely together is diminished, as analyzed in sections II and III above. Additionally, the intermediates may accumulate above the saturation threshold of the 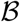 catalysts, at which point the efficiency of the second reaction becomes independent of the catalyst positions. To explore the combined effect of saturation and reduced intermediate leakage in our model, we adopt the partially absorbing boundary condition of Eq. S1 and adjust the leakage parameter *β* of Eq. S2. As seen in Fig. S9, this shifts the crossover point between the cluster and pair strategies back to smaller a values and decreases the enhancement achieved by both the clustering and the pair strategy.

**FIG. S9:**
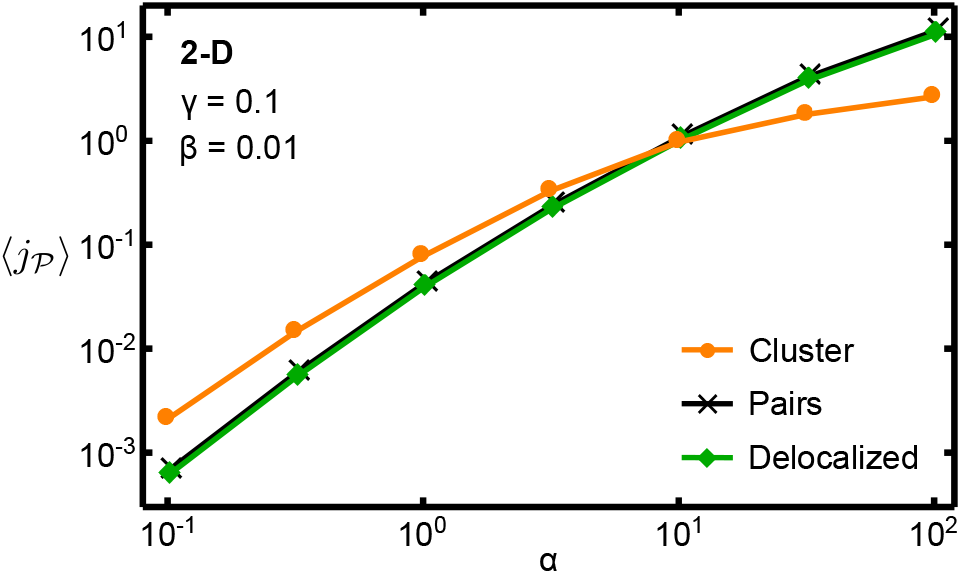
Catalyst saturation at reduced intermediate leakage. Comparison of the fluxes for delocalized, paired, and clustered catalysts for 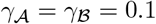 and *β* = 0.01.

In summary, this analysis shows that catalyst saturation does not strongly alter the qualitative behavior of the reaction fluxes of the different localization strategies. In large parts of the parameter space, we still find a crossover between the fluxes of the cluster and pair strategy as the reaction is changed from reaction-limited to diffusion-limited. However, the nonlinear reaction kinetics shifts the transition point to larger *α* values, due to the reduced impact of substrate depletion on the flux when the 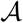 catalysts are saturated.

### VI. IMPACT OF STERIC EXCLUSION

To quantify steric exclusion effects, we calculated how the reaction flux is altered when the catalysts are made permeable to metabolites (Fig. 4 of the main text). For such a system, the steady-state concentration profiles of the substrate, 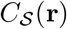, and the intermediate product, 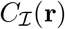, satisfy the reaction-diffusion balance equations

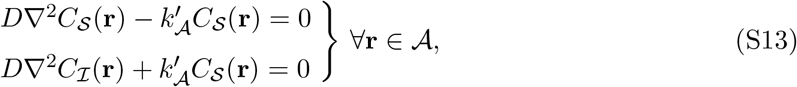

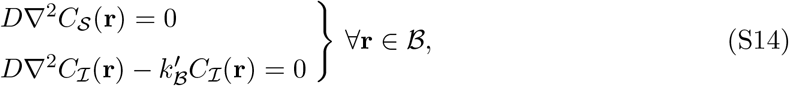

inside the volumes of the catalysts 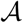 and 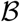, respectively. Outside of the catalyst volumes, 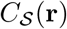 and 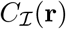 are stationary diffusion profiles satisfying

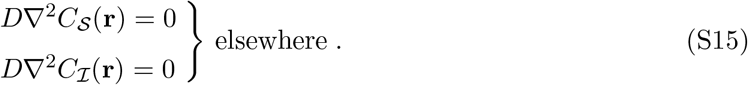

The catalytic activities 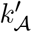 and 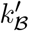 in Eqs. (S13) and (S14) are determined below as a function of the catalytic activities 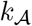 and 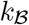 of the impermeable catalysts. Eqs. (S13–S15) are coupled via boundary conditions that require the concentration profiles to be continuous and smooth at the surfaces of all catalysts.

In Eqs. (S13) and (S14), the catalytic activity of the permeable catalysts 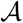 and 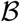 is homogeneously distributed over their volume, whereas the impermeable catalysts 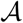 and 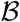 of the main text are catalytically active on their surface. To make the two models comparable, we have to choose the volume activities 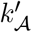 and 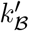 such that they produce the same overall catalytic activities as the surface activities 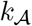 and 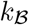. To this end we consider a system with a single permeable or impermeable catalyst and demand that the respective reaction fluxes are equivalent. To determine the reaction flux of a single 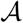 catalyst positioned at the system center, we solve the steady state diffusion equation, 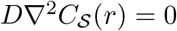, with the boundary conditions used in the main text,

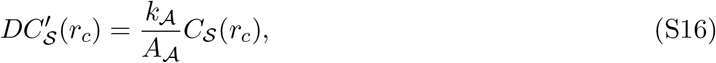

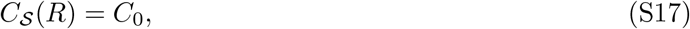

where *R* = 100 *r_c_* is the radius of the outer boundary. By solving this system, we obtain for the fluxes of an impermeable catalyst in 2 and 3 dimensions,

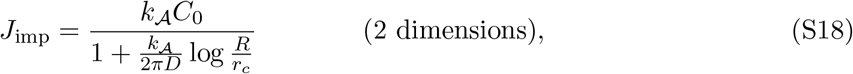

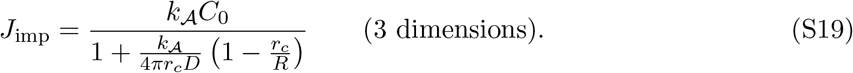

**FIG. S10:**
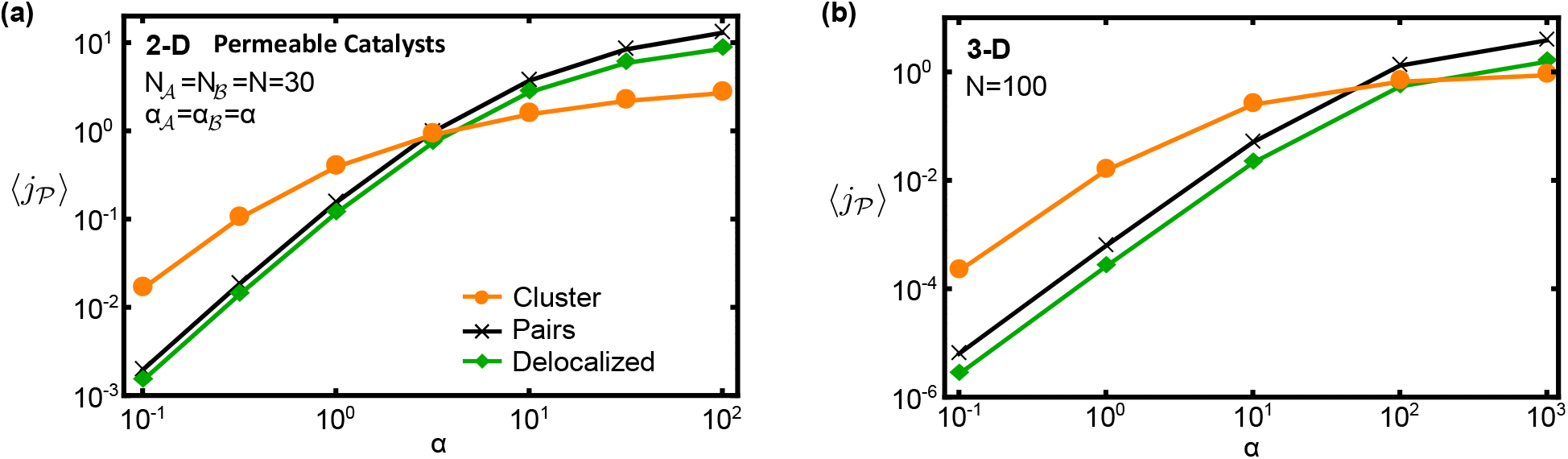
As in Figs. 3a and 3b of the main text, mean pathway fluxes of clustered arrangements, 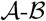-pair arrangements, and completely random arrangements (‘delocalized’) are shown as a function of the reaction-diffusion parameter *α*, but for permeable catalysts rather than impermeable catalysts. (a) Two-dimensional and (b) three-dimensional systems.

When a single central 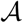 catalyst is permeable to substrate, the reaction flux can be computed by dividing the system into two domains, inside and outside of the catalyst, and solving in each domain the corresponding reaction-diffusion equation,

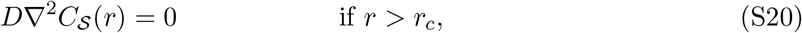

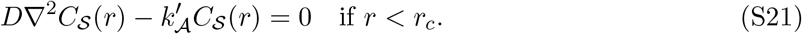

By requiring that the substrate concentration profile is continuous and differentiable at *r* = *r_c_*, together with the boundary conditions 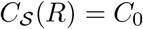 and 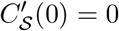, we can determine 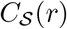. With this we compute the reaction flux by integrating the last term in Eq. (S21) over the domain of the catalyst. This yields the following expressions for the reaction flux of a permeable catalyst in 2 and 3 dimensions,

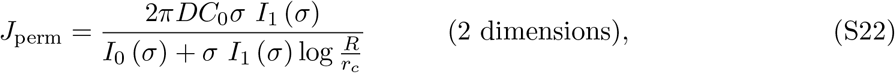

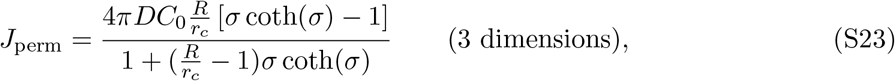

where 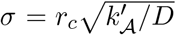 and *I_n_*(*x*) is the modified Bessel function of the first kind. We then choose 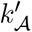 (as a function of 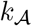) such that the fluxes *J*_perm_ and *J*_imp_ match. Comparing Eqs. (S18–S19) with Eqs. (S22–S23), we find that the fluxes will be equal when

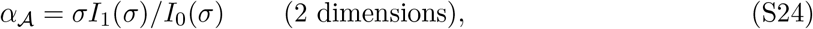

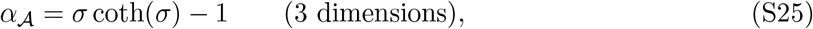

where 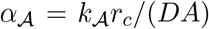 is the dimensionless reaction-diffusion parameter used in the main text (with *A* the surface area of the catalyst).

**FIG. S11:**
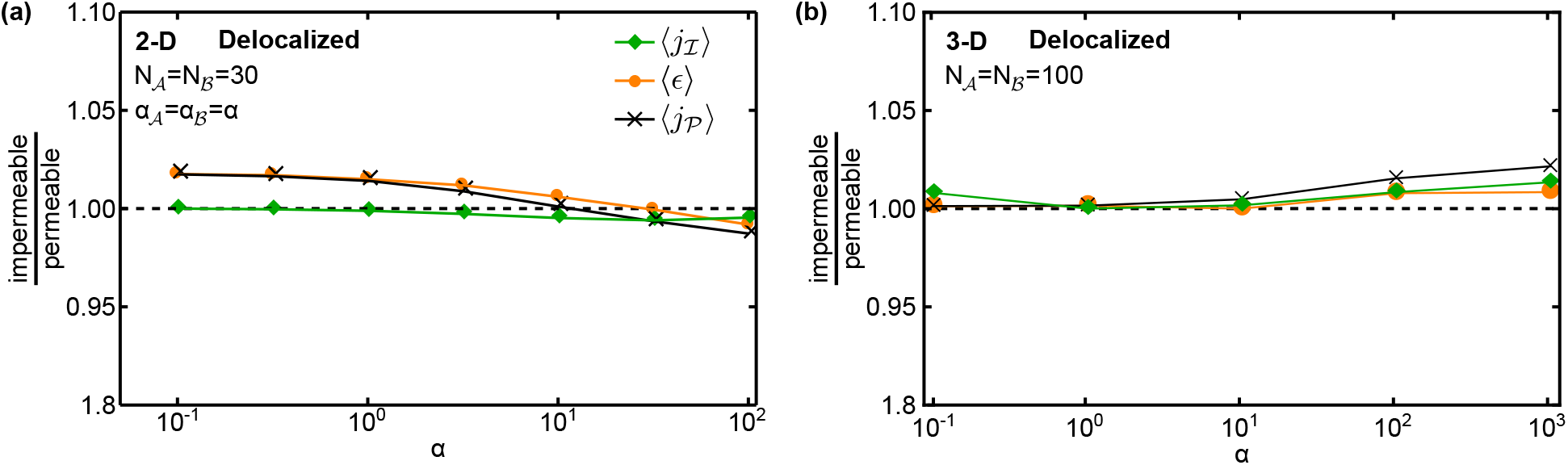
Comparison of 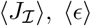, and 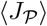 for impermeable versus permeable catalysts for delocalized arrangements in (a) two and (b) three dimensions.

We used the mappings (S24-S25) to determine the rate 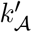 as a function of a 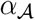 and the analogous mappings to determine the 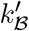 as a function of 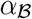. We then generated data for permeable catalysts by solving Eqs. (S13) to (S15) for the same catalyst configurations as with the impermeable catalysts; see Figs. S10a,b for permeable catalyst analogs of Figs. 3a,b in the main text. For catalyst configurations where steric exclusion effects are expected to be negligible, for example in the case of delocalized catalyst arrangements, this calculation led to reaction fluxes which are approximately the same for impermeable and permeable catalysts (see Fig. S11).

### VII. CATALYSTS WITH ACTIVE SITES

Molecular biocatalysts like enzymes are typically not reactive over their entire surface but rather have specific active sites at which reactions occur. To study the effect of such an anisotropy, we introduced model catalysts with a reactive patch covering 1/6 of the catalyst surface. The boundary conditions, Eqs. 5 and 6 of the main text, then only apply to the reactive fraction of the surface, while a no-flux boundary condition is applied on the remaining surface. To make these anisotropic model catalysts comparable to their isotropic counterparts, we compare them at the same total catalytic efficiency integrated over the respective surface area.

We first studied the pathway flux of anisotropic catalysts arranged according to the different localization strategies considered in the main text. Using ensembles of delocalized, paired, and randomly clustered arrangements, we computed the average pathway fluxes (Fig. S12) and compared them to the respective fluxes of isotropic catalysts (Fig. 3a). For small α values, the average flux of delocalized catalysts is not affected by the anisotropy (solid versus dashed green lines in Fig. S12). The same is true for randomly arranged clusters (solid versus dashed orange lines in Fig. S12). However, in both cases the average flux of anisotropic catalysts is slightly reduced compared to that of isotropic catalysts at larger α values (diffusion-limited regime). This is because the localization of the catalytic activity to a smaller region creates a stronger local depletion of metabolites around the reactive site, which in turn reduces the reaction flux. Nevertheless, the relative performance of the different strategies remains unchanged. For 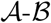 pair arrangements, we considered two relative orientations where the reactive patches either face each other (black line in Fig. S12) or point away from each other (blue line in Fig. S12). Comparing these two orientations, the pairs with facing sites produce a higher average flux over the entire α range.

We then determined the optimal configurations for complexes consisting of anisotropic catalysts. As in the main text (Fig. 5), we kept 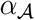 fixed and varied 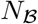 and 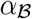 to explore the behavior of the optimal geometries and reactive site orientations. The anisotropy of the catalysts clearly changes the symmetries of the optimal arrangements (Fig. S13). While new geometries emerge that allow the reactive sites to be oriented optimally with respect to each other, the optimal configurations still appear to be determined by the trade-off between substrate shielding and intermediate confinement. In particular, when 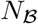 and 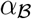 are varied, the optimal configurations display similar general trends as already observed in Fig. 5. For instance, the complexes are tightly packed at small 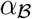 and open up as 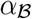 is increased. In the latter regime, some of the 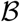 catalysts are localized close to 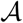, while others are positioned further away, at angular positions that cover “gaps” in the inner layer.

**FIG. S12:**
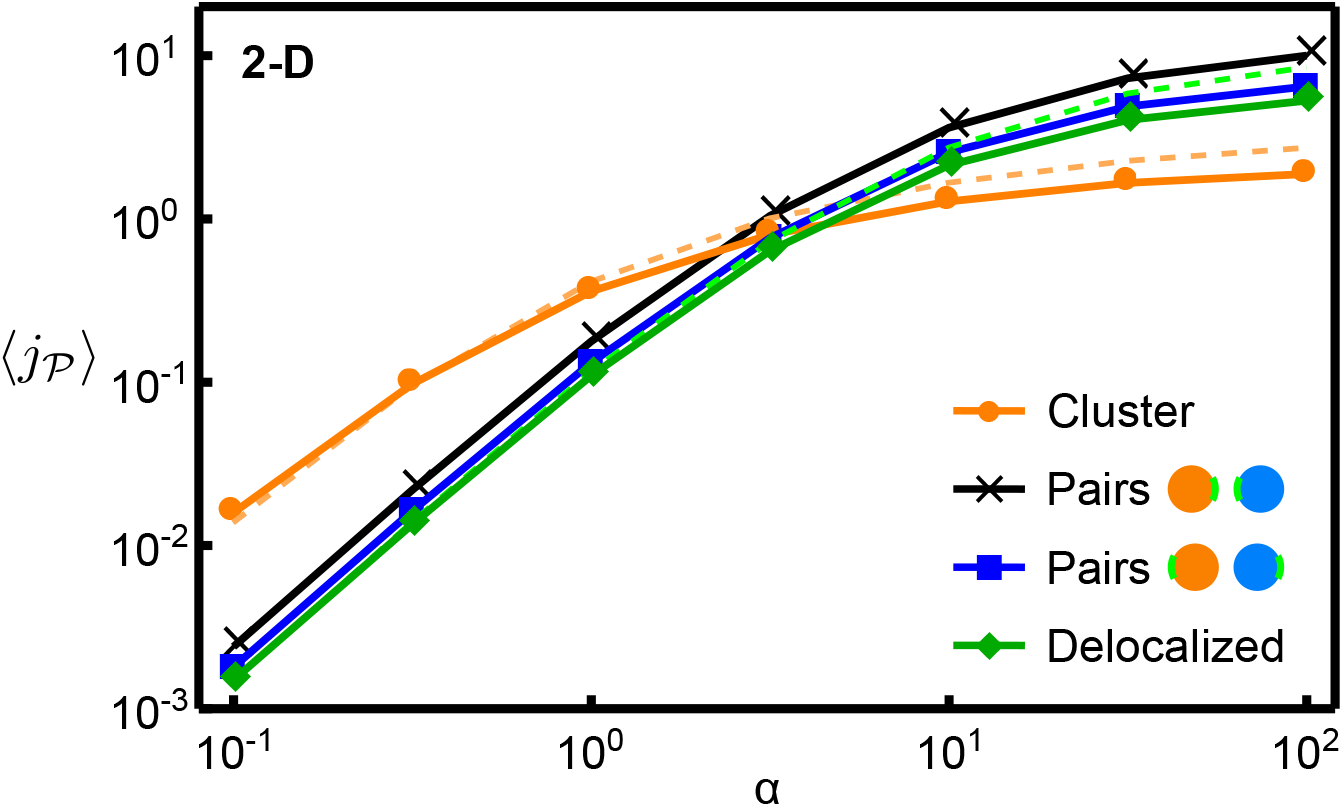
Comparison of the fluxes of delocalized, paired, and clustered catalysts with a reactive patch. The black and blue curve represents the reaction fluxes of the pair strategy with the reactive regions facing each other (black) and facing away from each other (blue). The dashed lines show the fluxes of the clustered and delocalized arrangements with uniform reactivity on the entire catalyst surface (Fig. 3a).

**FIG. S13:**
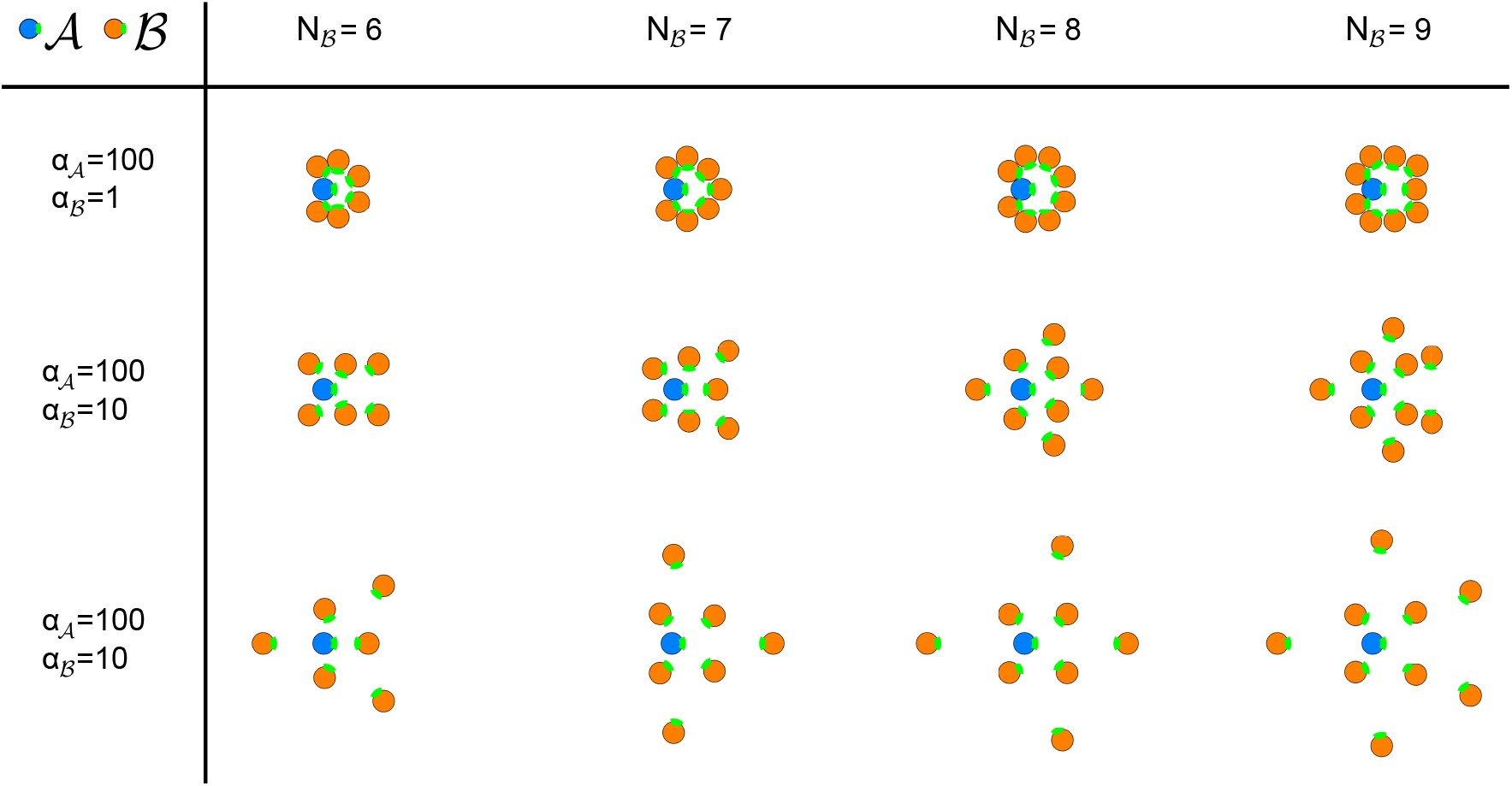
Optimal arrangements of several 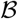 catalysts around a single 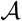 catalyst with reactive patches (green) comprising 1/6 of the catalyst surface.

### VIII. ENZYMES ARRANGED ON STRINGS AND SHEETS

An experimental study that is particularly interesting in the context of our theoretical analysis of different localization strategies was presented in Ref. [4]. This study arranged the same two consecutive enzymes in three different ways with the help of RNA scaffolds. Importantly, these experiments also included a loss mechanism for the intermediate product. Specifically, the two consecutive enzymes acyl-ACP reductase and aldehyde deformylating oxygenase were assembled with three different RNA scaffolds, and the conversion of hexadecanoyl-ACP to pentadecane was measured in the presence of competing enzymes that consume the intermediate product hexadecanal. The first scaffold colocalized the enzymes into pairs, the second scaffold arranged the enzymes into one-dimensional strings, and the third scaffold positions the enzymes on two-dimensional sheets.

To be able to relate our theoretical analysis to the experiments of Ref. [4], we also implemented the string and sheet arrangements using our 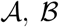 model catalysts, and computed the associated pathway fluxes. We implemented enzyme strings by placing alternating 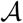 and 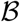 catalysts along a line in a 3D system. Similarly, we implemented enzyme sheets by placing 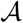 and 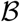 catalysts alternately on a planar square lattice, such that each 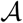 had four 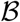 as nearest neighbors and *vice versa.* In both cases, we chose the center-to-center distance between neighboring catalysts to be 3*r_c_*, the same separation as for the pair arrangements analyzed in Fig. 3b of the main text. All other model assumptions for the enzyme strings and sheets were also the same as for the pairs in Fig. 3b.

The average pathway fluxes of the three different types of spatial arrangement are plotted in Fig. S14 as a function of the reaction diffusion parameter *α*, together with the reference curve for random delocalized arrangements. Fig. 6b of the main text additionally displays the average pathway flux for the clustered arrangements, showing that the catalyst sheets behave qualitatively similar to the random clusters: Just like the clusters, the sheets achieve a higher flux than the pairs in the reaction-limited regime (small *α*), whereas the pairs perform better in the diffusion-limited regime (large *α*). Steric effects can be expected to have a negligible influence on the flux generated by the sheet arrangement, because the access to the catalyst is not significantly blocked in such arrangements. Therefore, the observed behavior must be dominated by the effects of efficient intermediate transfer and substrate depletion.

In the diffusion-limited regime, the highest flux is achieved by the string arrangement. It shows a qualitatively similar behavior to the pair strategy, but displays a moderate overall enhancement in the flux. The string arrangement attenuates the detrimental effect of substrate depletion relative to the sheet and random cluster arrangements, while achieving an efficiency of intermediate processing that, although less than those of the cluster and sheet, is greater than in the pair arrangement.

**FIG. S14:**
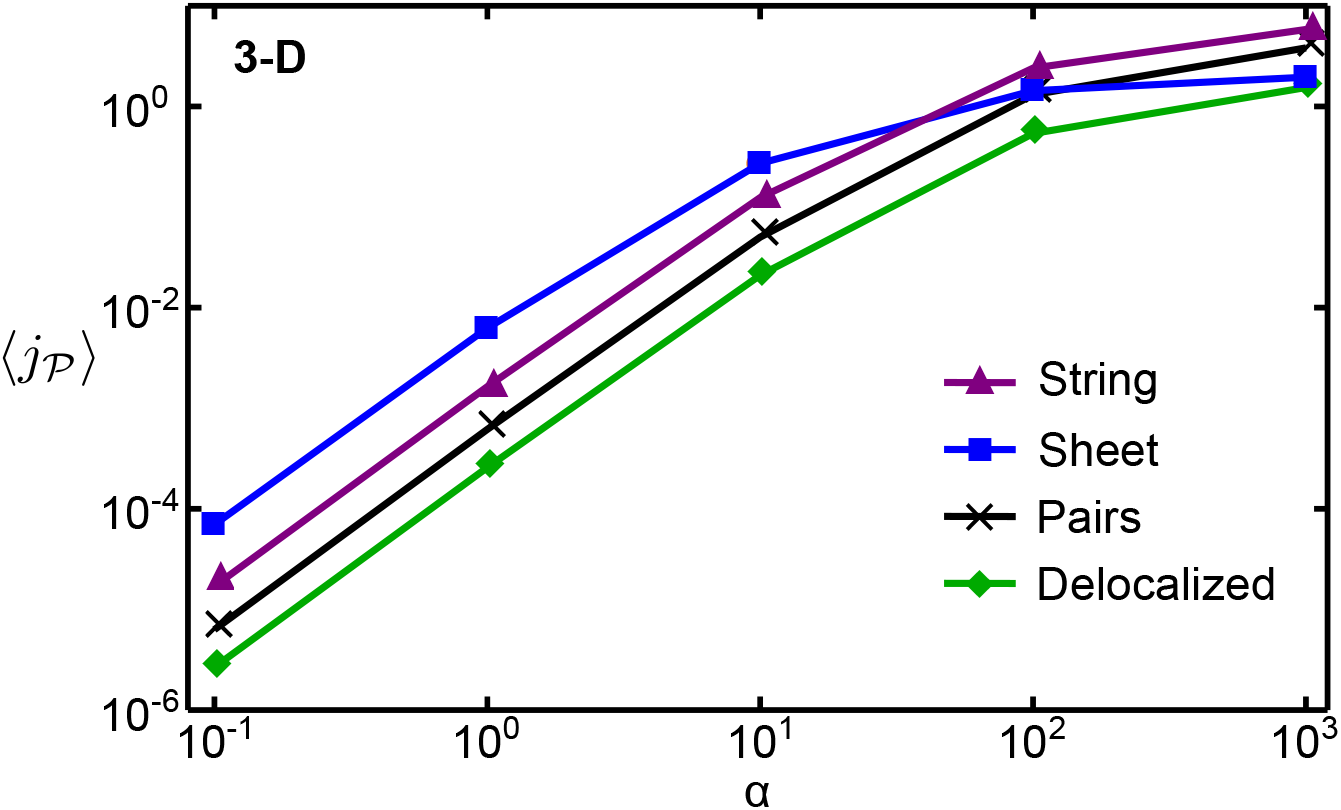
Comparison of the pathway fluxes of catalyst pairs, catalyst strings, and catalyst sheets, with the average flux of randomly delocalized catalysts shown as a reference. See text for parameters and model assumptions.

The consecutive enzymes used in the experimental study of Ref. [4] both lie in the reaction-limited regime, where our model displays the largest enhancement for the sheet arrangement, followed by a considerable enhancement for the string arrangement, and a small enhancement for the pair arrangement, all relative to delocalized catalysts. The *in vivo* measurements of pentadecane production with the three different RNA scaffolds display the same qualitative behavior [4], suggesting that our coarse-grained catalyst model can indeed capture the most essential features of engineered enzymatic systems.

## References

[1] Torquato, S. & Stillinger, F. H. Jammed hard-particle packings: From Kepler to Bernal and beyond. Rev. Mod. Phys. 82, 2633 (2010).

[2] Agapakis, C. M., Boyle, P. M. & Silver, P. A. Natural strategies for the spatial optimization of metabolism in synthetic biology. Nat. Chem. Biol. 8, 527–535 (2012).

[3] Srere, P. A. Complexes of sequential metabolic enzymes. Ann. Rev. Biochem. 56, 89–124 (1987).

[4] Jørgensen, K. et al. Metabolon formation and metabolic channeling in the biosynthesis of plant natural products. Curr. Opin. Plant Biol. 8, 280–291 (2005).

[5] Schmitt, D. L. & An, S. Spatial organization of metabolic enzyme complexes in cells. Biochemistry 56, 3184–3196 (2017).

[6] Kerfeld, C. A., Heinhorst, S. & Cannon, G. C. Bacterial microcompartments. Ann. Rev. Microbiol. 64, 391–408 (2010).

[7] Fontes, C. M. & Gilbert, H. J. Cellulosomes: highly efficient nanomachines designed to deconstruct plant cell wall complex carbohydrates. Ann. Rev. Biochem. 79, 655–681 (2010).

[8] An, S., Kumar, R., Sheets, E. D. & Benkovic, S. J. Reversible compartmentalization of de novo purine biosynthetic complexes in living cells. Science 320, 103–106 (2008).

[9] Yeates, T. O., Kerfeld, C. A., Heinhorst, S., Cannon, G. C. & Shively, J. M. Protein-based organelles in bacteria: carboxysomes and related microcompartments. Nat. Rev. Microbiol. 6, 681–691 (2008).

[10] Wheeldon, I. et al. Substrate channelling as an approach to cascade reactions. Nat. Chem. 8, 299–309 (2016).

[11] Lee, H., DeLoache, W. C. & Dueber, J. E. Spatial organization of enzymes for metabolic engineering. Metab. Eng. 14, 242–251 (2012).

[12] Küchler, A., Yoshimoto, M., Luginbühl, S., Mavelli, F. & Walde, P. Enzymatic reactions in confined environments. Nat. Nanotech. 11, 409–420 (2016).

[13] Lin, J.-L., Palomec, L. & Wheeldon, I. Design and analysis of enhanced catalysis in scaffolded multienzyme cascade reactions. ACS Catal. 4, 505–511 (2014).

[14] Yamada, Y. et al. Nanocrystal bilayer for tandem catalysis. Nat. Chem. 3, 372–376 (2011).

[15] Xie, C. et al. Tandem catalysis for CO_2_ hydrogenation to C_2_-C_4_ hydrocarbons. Nano Lett. 17, 3798–3802 (2017).

[16] Su, J. et al. Insights into the mechanism of tandem alkene hydroformylation over a nanostructured catalyst with multiple interfaces. J. Am. Chem. Soc. 138, 11568–11574 (2016).

[17] Müller, J. & Niemeyer, C. M. DNA-directed assembly of artificial multienzyme complexes. Biochem. Biophys. Res. Comm. 377, 62–67 (2008).

[18] Wilner, O. I. et al. Enzyme cascades activated on topologically programmed DNA scaffolds. Nat. Nanotech. 4, 249 (2009).

[19] Fu, J., Liu, M., Liu, Y., Woodbury, N. W. & Yan, H. Interenzyme substrate diffusion for an enzyme cascade organized on spatially addressable DNA nanostructures. J. Am. Chem. Soc. 134, 5516–5519 (2012).

[20] Zhang, Y., Tsitkov, S. & Hess, H. Proximity does not contribute to activity enhancement in the glucose oxidase–horseradish peroxidase cascade. Nat. Comm. 7, 13982 (2016).

[21] Wanders, R. J., Waterham, H. R. & Ferdinandusse, S. Metabolic interplay between peroxisomes and other subcellular organelles including mitochondria and the endoplasmic reticulum. Front. Cell Dev. Biol. 3, 83 (2015).

[22] Linka, M. & Weber, A. P. Shuffling ammonia between mitochondria and plastids during photorespiration. Trends Plant Sci. 10, 461–465 (2005).

[23] French, J. B. et al. Spatial colocalization and functional link of purinosomes with mitochondria. Science 351, 733–737 (2016).

[24] Ma, E. H. & Jones, R. G. TORCing up purine biosynthesis. Science 351, 670–671 (2016).

[25] Boetius, A. et al. A marine microbial consortium apparently mediating anaerobic oxidation of methane. Nature 407, 623–626 (2000).

[26] Müller, J. & Overmann, J. Close interspecies interactions between prokaryotes from sulfureous environments. Front. Microbiol. 2, 146 (2011).

[27] Mobarry, B. K., Wagner, M., Urbain, V., Rittmann, B. E. & Stahl, D. A. Phylogenetic probes for analyzing abundance and spatial organization of nitrifying bacteria. App. Env. Microbiol. 62, 2156–2162 (1996).

[28] Flores, E. & Herrero, A. Compartmentalized function through cell differentiation in filamentous cyanobacteria. Nat. Rev. Microbiol. 8, 39–50 (2010).

[29] Costa, E., Pérez, J. & Kreft, J.-U. Why is metabolic labour divided in nitrification? Trends Microbiol. 14, 213–219 (2006).

[30] Schramm, A. et al. Structure and function of a nitrifying biofilm as determined by in situ hybridization and the use of microelectrodes. App. Env. Microbiol. 62, 4641–4647 (1996).

[31] Buchner, A., Tostevin, F. & Gerland, U. Clustering and optimal arrangement of enzymes in reaction-diffusion systems. Phys. Rev. Lett. 110, 208104 (2013).

[32] Buchner, A., Tostevin, F., Hinzpeter, F. & Gerland, U. Optimization of collective enzyme activity via spatial localization. J. Chem. Phys. 139, 135101 (2013).

[33] Castellana, M. et al. Enzyme clustering accelerates processing of intermediates through metabolic channeling. Nat. Biotech. 32, 1011–1018 (2014).

[34] Minton, A. P. The influence of macromolecular crowding and macromolecular confinement on biochemical reactions in physiological media. J. Biol. Chem. 276, 10577–10580 (2001).

[35] Berry, H. Monte Carlo simulations of enzyme reactions in two dimensions: fractal kinetics and spatial segregation. Biophys. J. 83, 1891–1901 (2002).

[36] Schnell, S. & Turner, T. Reaction kinetics in intracellular environments with macromolecular crowding: simulations and rate laws. Prog. Biophys. Mol. Biol. 85, 235–260 (2004).

[37] Hu, Z., Jiang, J. & Rajagopalan, R. Effects of macromolecular crowding on biochemical reaction equilibria: a molecular thermodynamic perspective. Biophys. J. 93, 1464–1473 (2007).

[38] Dix, J. A. & Verkman, A. Crowding effects on diffusion in solutions and cells. Annu. Rev. Biophys. 37, 247–263 (2008).

[39] Nakamura, S., Hayashi, S. & Koga, K. Effect of periodate oxidation on the structure and properties of glucose oxidase. Biochim. Biophys. Acta Enzymol. 445, 294–308 (1976).

[40] Ban, Y. & Rizzolo, L. J. A culture model of development reveals multiple properties of rpe tight junctions. Mol. Vis. 3, 1 (1997).

[41] Pappenheimer, J., Renkin, E. & Borrero, L. Filtration, diffusion and molecular sieving through peripheral capillary membranes: a contribution to the pore theory of capillary permeability. Am. J. Physiol. 167, 13–46 (1951).

[42] Fogolari, F. et al. Studying interactions by molecular dynamics simulations at high concentration. BioMed Res. Internat. 2012 (2012).

[43] Martens-Habbena, W., Berube, P. M., Urakawa, H., José, R. & Stahl, D. A. Ammonia oxidation kinetics determine niche separation of nitrifying archaea and bacteria. Nature 461, 976–979 (2009).

[44] Albe, K. R., Butler, M. H. & Wright, B. E. Cellular concentrations of enzymes and their substrates. J. Theor. Biol. 143, 163–195 (1990).

[45] Samson, R. & Deutch, J. Exact solution for the diffusion controlled rate into a pair of reacting sinks. J. Chem. Phys. 67, 847 (1999).

[46] Tsao, H.-K. Competitive diffusion into two reactive spheres of different reactivity and size. Phys. Rev. E 66, 011108 (2002).

[47] Eun, C., Kekenes-Huskey, P. M. & McCammon, J. A. Influence of neighboring reactive particles on diffusion-limited reactions. J. Chem. Phys. 139, 044117 (2013).

[48] Hirakawa, H., Haga, T. & Nagamune, T. Artificial protein complexes for biocatalysis. Topics in Catalysis 55, 1124–1137 (2012).

[49] Shi, J. et al. Bioinspired construction of multi-enzyme catalytic systems. Chemical Society Reviews 47, 4295–4313 (2018).

[50] Liu, M. et al. A three-enzyme pathway with an optimised geometric arrangement to facilitate substrate transfer. ChemBioChem 17, 1097–1101 (2016).

[51] Reed, L. J. Multienzyme complexes. Acc. Chem. Res. 7, 40–46 (1974).

[52] Thomson, J. J. On the Structure of the Atom: an Investigation of the Stability and Periods of Oscillation of a number of Corpuscles arranged at equal intervals around the Circumference of a Circle; with Application of the Results to the Theory of Atomic Structure. Philos. Mag. 7, 237–265 (1904).

[53] Sachdeva, G., Garg, A., Godding, D., Way, J. C. & Silver, P. A. In vivo co-localization of enzymes on rna scaffolds increases metabolic production in a geometrically dependent manner. Nucleic Acids Res. 42, 9493–9503 (2014).

[54] Grotzky, A., Nauser, T., Erdogan, H., Schluter, A. D. & Walde, P. A fluorescently labeled dendronized polymer–enzyme conjugate carrying multiple copies of two different types of active enzymes. J. Am. Chem. Soc. 134, 11392–11395 (2012).

[55] Zhang, Y. et al. Using unnatural protein fusions to engineer resveratrol biosynthesis in yeast and mammalian cells. J. Am. Chem. Soc. 128, 13030–13031 (2006).

[56] Dueber, J. E. et al. Synthetic protein scaffolds provide modular control over metabolic flux. Nat. Biotech. 27, 753–759 (2009).

[57] Schwartz, R. E. The five-electron case of Thomson’s problem. Exp. Math. 22, 157–186 (2013).

[58] Yudin, V. The minimum of potential energy of a system of point charges. Discrete Math. App. 3, 75–82 (1993).

[59] Andreev, N. N. An extremal property of the icosahedron. East J. Approx. 2, 459–462 (1996).

[60] Smale, S. Mathematical problems for the next century. Math. Intell. 20, 7–15 (1998).

[61] Caspar, D. L. & Klug, A. Physical principles in the construction of regular viruses. In Cold Spring Harbor symposia on quantitative biology, vol. 27, 1–24 (Cold Spring Harbor Laboratory Press, 1962).

[62] Tarnai, T., Gaspar, Z. & Szalai, L. Pentagon packing models for “all-pentamer” virus structures. Biophys. J. 69, 612–618 (1995).

[63] Mackenzie, D. Proving the perfection of the honeycomb. Science 285, 1338–1339 (1999).

[64] Maritan, A., Micheletti, C., Trovato, A. & Banavar, J. R. Optimal shapes of compact strings. Nature 406, 287–290 (2000).

[65] Zandi, R., Reguera, D., Bruinsma, R. F., Gelbart, W. M. & Rudnick, J. Origin of icosahedral symmetry in viruses. Proc. Natl Acad. Sci. USA 101, 15556–15560 (2004).

[66] Nisoli, C., Gabor, N. M., Lammert, P. E., Maynard, J. & Crespi, V. H. Annealing a magnetic cactus into phyllotaxis. Phys. Rev. E 81, 046107 (2010).

[67] Agmon, N. & Szabo, A. Theory of reversible diffusion-influenced reactions. J. Chem. Phys. 92, 5270 (1990).

[68] Vijaykumar, A., Bolhuis, P. G. & ten Wolde, P. R. The intrinsic rate constants in diffusion-influenced reactions. Farad. Discuss. 195, 421–441 (2017).

[69] Vijaykumar, A., ten Wolde, P. R. & Bolhuis, P. G. The magnitude of the intrinsic rate constant: How deep can association reactions be in the diffusion limited regime? J. Chem. Phys. 147, 184108 (2017). URL https://doi.org/10.1063/1.5009547.

## References

[1] Fu, J., Liu, M., Liu, Y., Woodbury, N. W. & Yan, H. Interenzyme substrate diffusion for an enzyme cascade organized on spatially addressable DNA nanostructures. J. Am. Chem. Soc. 134, 5516–5519 (2012).

[2] Wheeldon, I. et al. Substrate channelling as an approach to cascade reactions. Nat. Chem. 8, 299–309 (2016).

[3] Zhang, Y., Tsitkov, S. & Hess, H. Proximity does not contribute to activity enhancement in the glucose oxidase–horseradish peroxidase cascade. Nat. Comm. 7, 13982 (2016).

[4] Sachdeva, G., Garg, A., Godding, D., Way, J. C. & Silver, P. A. In vivo co-localization of enzymes on rna scaffolds increases metabolic production in a geometrically dependent manner. Nucleic Acids Res. 42, 9493–9503 (2014).

